# Distinct neurophysiological signatures of guided visual search in schizophrenia and autism: Evidence from fixation-related potentials, alpha modulation, and functional MRI

**DOI:** 10.64898/2026.11.17.767357

**Authors:** Eduardo Gonzalez-Moreira, Antigona Martinez, Maria B. Aburto-Ponce, Maxwell Greisberg, Odeta Beggel, Russell H. Tobe, Alexandre R. Franco, Elisa Dias, Daniel C. Javitt

**Author notes:** **Correspondence:** Eduardo Gonzalez-Moreira.

## Abstract

Schizophrenia (SZ) and autism spectrum disorder (ASD) are associated with convergent differences in social perception relative to non-autistic, non-schizophrenia controls (NC), but divergent patterns of alterations in early visual activity. Here, we used multimodal imaging during the Guided Visual Search task to investigate the differential patterns of early visual alteration across SZ and ASD including superior colliculus (SC) and pulvinar nucleus. In Experiment 1, simultaneous eye tracking and EEG were obtained from 91 individuals (35 NC, 20 ASD, 36 SZ). Fixation-related potential (FRP) was time-locked to fixation onset. In Experiment 2, fMRI during Guided Search was obtained from a partly overlapping sample of 71 individuals (27 NC, 25 ASD, 19 SZ). In behavioral analyses, SZ was associated with significantly prolonged search time (d=.74, p=.003), which was attributable primarily to reduced mean saccade amplitude, whereas ASD was associated with a small advantage (d=.35), which was not statistically significant. In EEG analyses, SZ was associated with reduced amplitude of the FRP P1 (lambda) response (d=1.15, p<.0001) relative to NC that remained significant following control for saccade size (p<.0001), but with normal task-related modulation of background alpha activity. In contrast, ASD was associated with normal FRP amplitude and larger alpha suppression (d=.83, p=.007) that was attributable to increased alpha levels during pre-task fixation (at matched pre-onset alpha levels, d=.14, p=.66). Within SZ, P1 correlated with the MCCB composite (r=.56, p=.001). Within ASD, ADOS-2 Communication correlated with P1 (r=.51, p=.024) and with the broadband change from pre-onset to search (r=−.51, p=.030). In fMRI, SZ was associated with correlated activation reductions in the Visual1 (Sidak p<.001) and Visual2 (Sidak p=.022) networks and SC (Sidak p=.019), whereas activation was unaffected in the Dorsal Attention and Frontoparietal networks, Frontal Eye Fields and pulvinar. In ASD, activation was increased in pulvinar (Sidak p≤.038), but unaffected elsewhere. Cortical activation patterns associated with SC activation resembled those associated with P1 generation across groups. Overall, the findings support bottom-up disturbances of visual processing in schizophrenia. In autism, pre-onset alpha was higher and task-related suppression was larger.

## 1 Introduction

Schizophrenia (SZ) and autism spectrum disorder (ASD) are highly prevalent mental health conditions associated with convergent reductions in visual aspects of social cognition, such as face emotion recognition (FER) [Kohler et al., 2010; Uljarevic & Hamilton, 2013; Ozbek et al., 2023] which have been linked to divergent patterns of early visual processing. Thus, in SZ, impairments in FER are linked to reduced activation in primary (V1) and secondary (V2) visual cortices as measured using fMRI, as well as reductions in activation in lateral and medial subdivisions of pulvinar nucleus of the thalamus. In contrast, in ASD reductions in FER have been linked to increased activation both of V2 and the inferior subdivision of pulvinar [Martinez et al., 2019; Martinez et al., 2021].

In electrophysiological studies, SZ is associated with reduced stimulus-driven sensory responses and stimulus-driven reductions in ongoing alpha power. In contrast, the stimulus-induced alpha suppression responses are enhanced in ASD to both simple visual stimuli [Martinez et al., 2019] and faces [Javitt et al., 2026], and in both cases the increases correlate strongly with difficulties in social interaction as reflected in the Autism Diagnostic Observation Schedule, Second Edition (ADOS-2) [Lord et al., 2012] social interaction score [Martinez et al., 2019; Javitt et al., 2026].

We have also previously observed that SZ is associated with reduced fixation-related potentials (FRP) during both simple visual search [Dias et al., 2020] and reading [Dias et al., 2021] paradigms and that these deficits correlate with impaired neurocognitive function. Here, in order to gain further insight into differential patterns of early visual processing in SZ and ASD, we evaluated patterns of behavioral, EEG and fMRI responses during the same search task that we previously studied in SZ alone.

We chose this task in part because it has been recommended as a paradigmatic task related to control of attention within the early visual system [Nuechterlein et al., 2009], and in part because prior literature suggests that SZ and autistic individuals show differentiable patterns of divergent function relative to non-autistic, non-schizophrenia controls (NC). Thus, SZ is associated with consistent, large effect-size (Cohen’s d∼.8) increases in search time during serial search [Dias et al., 2020; Nuechterlein et al., 2009; Sklar et al., 2020] relative to NC. In contrast, autism is associated with a small-moderate (Cohen’s d∼.25) advantage, dependent upon task parameters [Kaldy et al., 2016; Constable et al., 2020]. Theories of autistic advantage include both 1) perceptual enhancement and 2) hyper-focus during the task, leading to enhanced attentional engagement [Kaldy et al., 2016].

In our study, we assessed two different sets of functions. First, we collected combined eye tracking and continuous EEG data during guided search on a group of 91 participants including SZ, autistic and NC individuals. Second, we collected fMRI during guided search from 71 individuals, who partially overlapped the EEG cohort. Our analyses focused on across-group differences within modalities, as well as correlations across modalities. In both cases, we used a conjunction search task in which stimuli (Gabor patches) differed by both orientation and contrast, and participants needed to detect a single stimulus in each field with a pre-designated conjunction of features.

In the eye-tracking/EEG dataset, we evaluated total search time per field, mean saccade amplitude between fixations across sequential eye movements and fixation duration. In a previous study, we observed that the saccade immediately preceding target fixation (as opposed to other saccades) was associated with late delta activity, suggestive of post-fixation target detection within peripheral space, which then led to a saccade to the target. SZ was associated with normal late delta FRP response to the penultimate fixation, but the saccade amplitudes were themselves reduced, suggesting reduced ability to detect the targets within parafoveal space. The reduction in saccade amplitude correlated with increased search time. In contrast, fixation duration was not different between groups [Dias et al., 2020].

In addition, we evaluated two informative neurophysiological measures. First, we assessed amplitude of the FRP, which is obtained by averaging responses that are time locked to fixation onset. FRP are considered to contain contributions from both sensory driven activity [Dimigen & Ehinger, 2021; Kadosh & Bonneh, 2023; Snell et al., 2023; Del Campo et al., 2024] and from corollary discharge originating within the oculomotor system [Rajkai et al., 2008]. Second, we evaluated the suppression of alpha activity over occipital regions during the task, which is considered to be an index of task-engagement [Green et al., 2017; Pfurtscheller & Lopes da Silva, 1999; Klimesch, 2012]. In our study, FRP in SZ were significantly reduced in amplitude, supporting concepts of impaired bottom-up and corollary discharge-driven activity [Spering et al., 2013]. In time-frequency analyses [Javitt et al., 2020], the FRP-P1 response corresponded primarily to reduced modulation of intertrial coherence (ITC) in the theta (4-8 Hz) frequency range. In contrast, task-related alpha suppression was unaffected, suggesting intact top-down attentional modulation [Dias et al., 2021].

Here, in Experiment 1 we conducted a follow-up combined eye-tracking and FRP study in a parallel group of SZ and adult autistic individuals, with the goal of further examining the neural mechanisms of divergence across disorders. Although the serial search paradigm has been studied extensively in ASD, to the best of our knowledge this is the first study to collect neurophysiological biomarkers. Given the known autistic advantage, we hypothesized that ASD might be associated with either 1) increased FRP, which would potentially indicate increased sensitivity to visual input or corollary discharge, or 2) increased task-related alpha modulation, which might indicate task-related hyper-engagement.

In addition, we conducted a second experiment in which we collected fMRI concurrent with visual search in both SZ and autistic groups, relative to NC. At present, despite the well-replicated behavioral deficits in guided search in SZ, relatively little is known about the underlying substrates. At the cortical level, the guided search task engages multiple cortical networks, including visual sensory, dorsal attention and fronto-parietal “task positive” networks whereas at the subcortical level both pulvinar [Strumpf et al., 2013] and superior colliculus [Himmelbach et al., 2007] have been implicated [Kim et al., 2012]. Our goal therefore was to determine potential substrates of impaired visual search in SZ and potentially superior visual search in ASD at both the cortical and subcortical levels. Within the oculomotor system, both the frontal eye fields (FEF) and superior colliculus (SC) are potential sources of corollary discharge input into visual cortex. Specific analyses therefore focused on activation within these regions relative to FRP amplitude and overall activation patterns.

Consistent with prior literature, we hypothesized that SZ and autistic participants would show differential patterns of divergence with increased and reduced search times, respectively. In SZ, we hypothesized that impaired search performance would be associated with reduced activation within subcortical structures such as SC and pulvinar, along with early visual regions. In ASD, we hypothesized that we would see enhanced top-down modulation of alpha suppression analogous to the enhanced stimulus-driven suppression that we have observed in prior studies [Martinez et al., 2019; Martinez et al., 2021; Javitt et al., 2026], and that this would be beneficial for processing of the non-social stimuli used for the present study. Finally, across subjects, we hypothesized that FRP amplitude would correlate with SC activation levels across groups, consistent with a role of SC as a mediator of corollary discharge input into V1 and early visual cortex.

## 2 Materials and Methods

### 2.1 Participants

Participants were adult individuals of either sex diagnosed with ASD or SZ, or non-autistic, non-schizophrenia controls (NC), free of other major neuropsychiatric disorders (ages and other characteristics in Table 1). Diagnosis was made using the Structured Clinical Interview for DSM disorders [First et al., 2002; First et al., 2015]. All participants had normal or corrected-to-normal vision, 20/32 or better on the Logarithmic Visual Acuity Chart (Precision Vision, LaSalle, IL). Estimated premorbid IQ was assessed using the Ammons Quick Test [Ammons & Ammons, 1962]. Participant and parental socioeconomic status (SES) was assessed using the Hollingshead Four Factor Index of Social Status [Hollingshead, 2011]. In ASD, symptoms were assessed using the ADOS-2 [Lord et al., 2012]. In SZ, symptoms were assessed using the Positive and Negative Syndrome Scale (PANSS) [Kay et al., 1987], while cognition was assessed using the MATRICS Consensus Cognitive Battery (MCCB) [Nuechterlein et al., 2008].

**Table 1.**
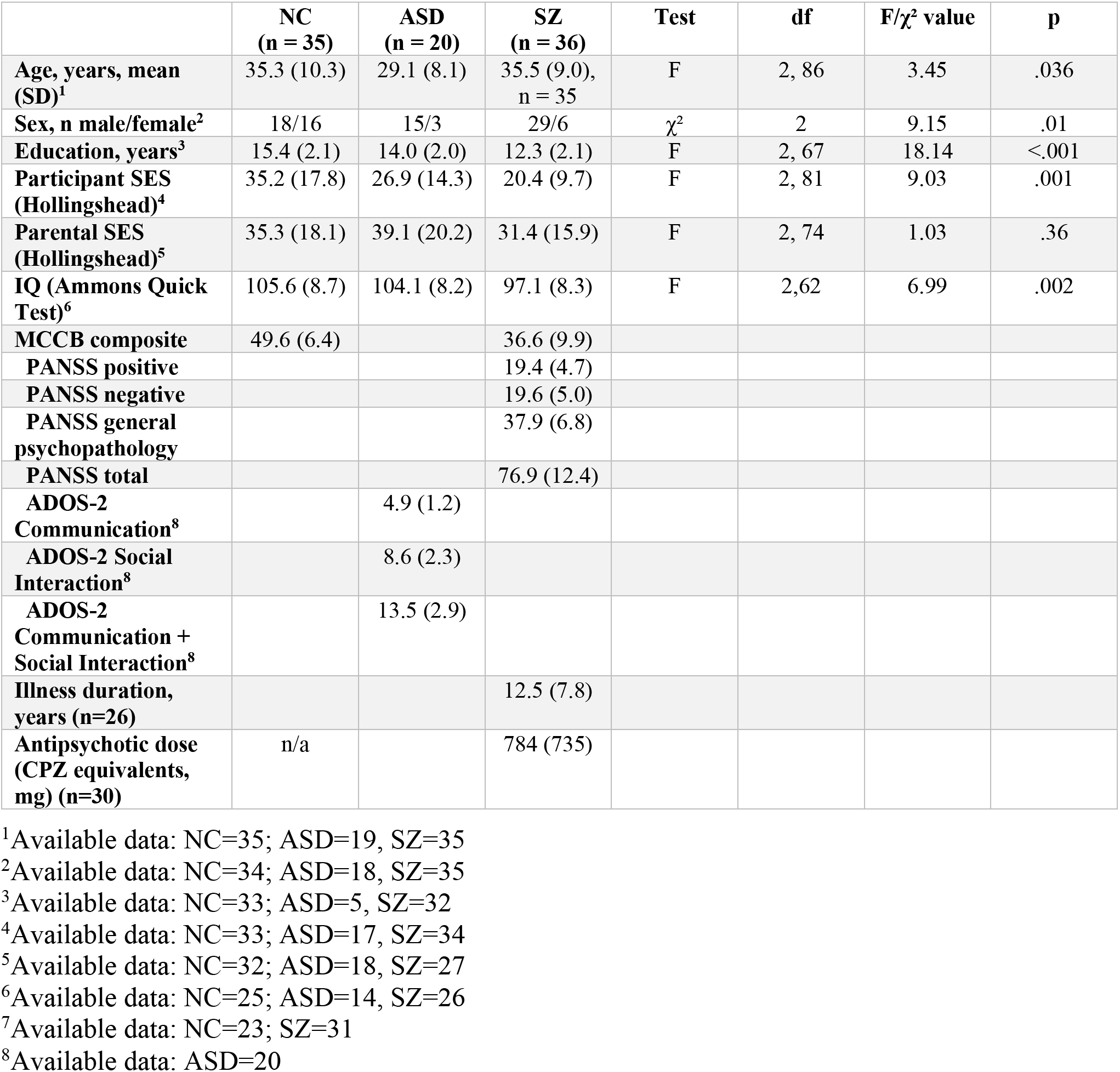
Participant characteristics. Values are mean (SD). Between group comparisons are performed using ANOVA or Chi-square for continuous and categorical data as appropriate.

The overall study contained separate EEG (Experiment 1) and MRI (Experiment 2) experiments. All procedures for both experiments were approved by the Institutional Review Board of the Nathan S. Kline Institute for Psychiatric Research and were conducted in accordance with the Declaration of Helsinki. All subjects provided written informed consent after receiving a complete description of the study and were compensated for their participation.

The Experiment 1 groups differed in age, sex, education, participant SES and estimated IQ (Table 1). Autistic participants were younger than NC. Both clinical groups included a higher proportion of men. Years of education, participant SES, and estimated IQ were lower in SZ; the education mean in autism is based on five participants and is not interpreted. Parental SES did not differ across groups. Within families, NC did not differ from their parents (parent minus participant, −1.3; t(30) = −0.53, p = .60), whereas autistic participants (+13.8; t(16) = 3.91, p = .001) and participants with SZ (+10.3; t(26) = 4.47, p < .001) scored lower than their parents (paired t-tests; higher Hollingshead scores indicate higher socioeconomic status).

### 2.2 Visual search task

Stimuli were conjunction search arrays of Gabor patches differing in both orientation and contrast, and each field contained a single target defined by a pre-designated conjunction of the two features, as in Dias et al. (2020). In the serial condition analyzed here, the target was a low-contrast, vertically oriented patch among high-contrast vertical and low-contrast horizontal distractors, so that neither contrast nor orientation alone identified it; all patches had the same spatial frequency. Each field contained 48 patches, half at high contrast and half at low (40%) contrast, at random positions within a virtual 8 by 8 grid. Each field was preceded by drift correction and a screen indicating the contrast of the upcoming target, and fields timed out after 30 s. Experiment 1 combined eye tracking and EEG during guided search: 91 participants contributed behavior and eye tracking (NC 35, ASD 20, SZ 36), and 90 contributed EEG after quality criteria (NC 35, ASD 20, SZ 35), in two cohorts of 86 that are not nested (fixation-related measures NC 34, ASD 19, SZ 33; display-locked alpha NC 33, ASD 18, SZ 35; 82 participants are in both). Experiment 2 investigated fMRI activation patterns in a partially overlapping sample (n=71; NC 27, ASD 25, SZ 19). Thirty-two participants had both usable EEG and usable fMRI (NC 5, ASD 16, SZ 11).

In Experiment 1, participants performed the serial visual search task in four blocks of 48 trials (192 trials in total). Each trial began with a central fixation, followed by a search display that remained visible until the participant located the target or the field timed out. As in Dias et al. (2020), search fields were 21° square and were presented on an Iiyama Vision Master Pro 512 CRT monitor (85 Hz, 1024 by 768 pixels) at a viewing distance of 57 cm, with the same set-up for all three groups. Search time was defined as the onset of the final fixation, so that dwell time on the target was excluded and the measure reflected the time taken to reach the target rather than to respond.

In Experiment 2, the same serial conjunction-search stimuli were adapted to a blocked fMRI design. Visual-search and fixation-control periods were modeled as 12-s blocks. During visual-search blocks, participants searched for the predefined target, whereas during fixation-control blocks they viewed comparable stimulus arrays while maintaining central fixation and responded when the fixation cross blinked. Intermittent 2-s response prompts asked participants to indicate whether the target had appeared in the left or right visual hemifield; instruction periods (4 s) and response prompts were modeled separately.

### 2.3 Experiment 1: behavior, eye tracking and EEG

#### 2.3.1 Behavior

The primary behavioral variable was search time, the onset of the final fixation in each search array, averaged across arrays. Search time was the onset of the last on-screen fixation in each trial. Fixations outside the 1024 × 768 display were removed, trials with fewer than three remaining fixations were excluded, and search time was averaged over trials within each block and then over blocks. For each search field, participants indicated that they had detected the target by pressing a button on a gamepad. As in Dias et al. (2020), a response was accepted as correct only if a fixation had been made within a 3° window around the target within 2 s of the button press; the same rule applied to all three groups.

#### 2.3.2 Eye tracking

Gaze was recorded with a remote EyeLink-1000 system (SR Research, Ottawa, Canada) at 500 Hz in pupil and corneal-reflection mode, monocularly from the right eye in all but four of the 350 serial-search recordings, with pupil size in area units. Participants were seated in a darkened, electrically shielded room with the head stabilized on a chinrest 57 cm from the monitor. A nine-point calibration and validation procedure (HV9) was used, and drift correction was applied before each trial while participants fixated a central dot. In the 174 serial-search recordings that contain a validation (223 validations), the median average error was 0.54 degrees (range 0.13 to 2.01) and the median maximum error 1.26 degrees (range 0.45 to 13.20); 93% of validations had an average error of one degree or less. The 13.20-degree value was a first validation that was repeated; the last validation in a recording did not exceed 4.05 degrees. Saccades and fixations were classified with the EyeLink cognitive parser configuration (motion threshold 0.1 degrees, velocity threshold 30 degrees/s, acceleration threshold 8,000 degrees/s²), and a fixation was defined as the interval following the end of the preceding saccade. These values were read from the raw recordings. Saccade amplitude in the Results is the distance between successive fixations; the tracker’s own per-saccade amplitude correlated only moderately with it (r =0.52) and was used as a check, on which the SZ–NC difference was not significant (see Supplementary Material).

#### 2.3.3 EEG acquisition and preprocessing

EEG was recorded during the search task with two amplifier systems: an ANT Waveguard and a BrainVision system, which were balanced across groups in the display-locked alpha cohort (ANT/BrainVision: NC 21/12, ASD 13/5, SZ 23/12; chi-square p=.82). Impedances were maintained below 10 kΩ. The ANT system sampled at 1024 Hz and the BrainVision system at 500 Hz. Data were processed offline in Python and resampled to a common 250 Hz. For the alpha measures, recordings were filtered from 1 to 45 Hz with a fourth-order zero-phase Butterworth filter and re-referenced to the common average, and epochs in which the absolute amplitude exceeded 350 µV were rejected (at any channel for the spectral measures, and at PO3, POz or PO4 for the time course). For the fixation-related measures, ocular artifacts were removed with an ICA procedure optimized for free viewing [Dimigen, 2020], which removed one to four blink-related components per participant, and the recordings were then filtered from 0.1 to 80 Hz with the same type of filter; the alpha measures were computed without ICA. These settings describe the analysis code applied to the stored recordings, not any earlier processing of those files. No separate resting recording was used: the pre-onset alpha level was measured in, and the 8–12 Hz change relative to, the second preceding display onset (−1 to 0 s) within the search task, and the display-locked alpha time course and the measures taken from it (sustained and peak suppression, latency to half-maximum, slope and time to peak suppression) against −0.6 to −0.2 s.

#### 2.3.4 Fixation-related potentials (FRP)

Fixation-related epochs were extracted from −600 to +600 ms around fixation onset; epochs with a peak-to-peak amplitude above 150 µV on any channel between −300 and +500 ms were rejected, and the remaining epochs were re-referenced to the common average. FRP epochs were baseline-corrected to −500 to −300 ms. The occipital region of interest comprised O1, Oz and O2. Blink-related components were removed with the ICA procedure described in 2.3.3. The P1 response amplitude was defined as the mean amplitude from 80 to 140 ms after fixation onset. Theta ITC was computed by Morlet decomposition over 4-7.5 Hz and quantified as the maximum ITC from 50 to 250 ms after fixation onset. Because ITC is biased upward at low trial counts, the ITC estimate used the first 600 accepted epochs of each participant: 600 for 83 participants, and 318 and 541 for two ASD participants with fewer accepted epochs. Excluding these two left the group difference significant (F(2,80)=13.17, p<.0001). Theta ITC required at least 40 accepted epochs, and the trial-level mixed model at least 200 accepted fixations. One SZ participant, with fixation reports from two of the four blocks and 36 accepted epochs, contributed to neither, so those two analyses include 85 participants (NC 34, ASD 19, SZ 32), whereas P1 and the per-participant regressions include 86.

Responses were quantified by averaging epochs, with overlap and saccade amplitude addressed by the isolated-fixation criterion and the trial-level mixed model.

#### 2.3.5 Task-related alpha suppression

Alpha measures were taken from PO3, POz, and PO4, parieto-occipital sites that match the known posterior scalp distribution of alpha and at which the change, collapsed across groups, was maximal, so that the choice does not depend on the group difference (Supplementary Material). Task-related suppression, the primary alpha measure, was the change from the pre-onset window (−1 to 0 s) to the search window (1 to 2 s) over the 8–12 Hz band for trials lasting at least 2.5 s. To quantify it, we computed the change ratio in decibels (dB), 10log10(search/pre-onset power), across the 5-25 Hz frequency range. Consistent with the calculation, suppression is expressed as a negative value.

The frequency resolution of both windows is 1 Hz, set by their length; zero-padding interpolates the plotted spectrum at 0.25 Hz intervals without increasing its effective resolution. The pre-onset window could not be lengthened because the median interval between the end of one trial and the start of the next was 1.06 s.

In order to separate periodic from aperiodic (1/f) alpha, we parameterized each participant’s pre-onset spectrum at PO3/POz/PO4 with fooof [Donoghue et al., 2020] (version 1.1.1, since renamed specparam; 2-30 Hz, fixed aperiodic mode, peak-width limits 1-8 Hz, at most four peaks), which fits the aperiodic component and the periodic peaks above it. Periodic alpha power was the height above the aperiodic fit of the largest peak between 7 and 14 Hz. Pre-onset alpha was also expressed as periodic power after removal of the aperiodic component; participants without a detected peak (n=7) were excluded from that measure.

For the scalp-distribution and individual-peak analyses, the pre-onset and search-window spectra were recomputed at all channels, using the same epochs, windows, trial-duration criterion and artifact threshold. Each participant’s pre-onset alpha peak was the periodic peak detected after removal of the aperiodic component from the PO3/POz/PO4 pre-onset spectrum (Supplementary Methods).

In addition to alpha amplitude, we evaluated time to peak alpha suppression, the time of the minimum of each participant’s display-locked 8–12 Hz time course between 0 and 2 s after display onset; the time course was Morlet power (4 cycles, 8 to 12 Hz in 1 Hz steps) at PO3/POz/PO4 in trials lasting at least 2.5 s (at least 15 per participant), averaged over channels, frequencies and trials and expressed in dB relative to −0.6 to −0.2 s.

### 2.4 fMRI

For fMRI, participants completed the visual-search task across two functional runs which were analyzed jointly. Task fMRI used a block design comprising 12-s visual-search and fixation-control blocks, with 2-s response-question periods and 4-s instruction periods modeled separately. Functional images were acquired on a 3-T Siemens MAGNETOM TrioTim scanner using a multiband gradient-echo EPI sequence (TR=1400 ms, TE=30 ms, 64 slices, 2-mm isotropic voxels, multiband acceleration factor=4, field of view=224 mm). High-resolution T1-weighted anatomical images were acquired using an MPRAGE sequence (TR=1900 ms, TE=2.52 ms, TI=900 ms, flip angle=9°, 176 sagittal slices, 1-mm isotropic voxels, field of view=250 mm).

Functional data were processed and analyzed in AFNI using afni_proc.py [Reynolds et al., 2024]. Preprocessing included slice-timing correction, alignment of the functional data to the anatomical image, spatial normalization to the MNI152 2009 template at 2-mm isotropic resolution, motion correction, spatial smoothing with a 4-mm FWHM Gaussian kernel, and voxelwise scaling. Time points exceeding a motion threshold of 0.3 mm were censored. Demeaned motion parameters and their temporal derivatives were included as nuisance regressors, with motion regressors modeled separately for each run. Data from both runs were modeled jointly at the individual-subject level using a general linear model. Visual-search and fixation-control periods were modeled as 12-s blocks, while response-question (2 s) and instruction (4 s) periods were modeled separately. Six 10-s rest blocks consisting of only a fixation cross were interspersed throughout the task but were not explicitly modeled.

Mean beta coefficients were extracted separately for visual-search (SEARCH) and fixation-control (FIX) conditions, as well as for their contrast (SEARCH–FIX). Cortical network-level beta coefficients were estimated using the 12-network cortical-subcortical partition of Ji and colleagues [Ji et al., 2019]. Because this partition does not provide anatomical definitions of the superior colliculus or pulvinar, these structures were defined using the atlas of McFadyen and colleagues [McFadyen et al., 2019], with masks collapsed across hemispheres.

### 2.5 Statistical analysis and exclusions

Group comparisons used one-way analysis of variance with Sidak post-hoc contrasts, and all three pairwise contrasts are reported with Cohen’s d. Associations were tested with Pearson correlations, both pooled and after centering each variable on its group mean. For behavior measures, the primary variable of interest was mean search time per field. Secondary measures included mean saccade amplitude for random and pre-final fixations and fixation duration.

For fMRI, the primary analyses focused on task-related activation (SEARCH-FIX) within cortical networks known to be engaged by the task (Visual1, Visual2, Dorsal Attention, Frontoparietal), as well as superior colliculus, pulvinar and FEF. Exploratory analyses evaluated between-group differences during the search and fixation blocks separately, relative to rest. Finally, brain-wide regressions against behavioral (search time) and FRP measures evaluated whether additional cortical regions were associated with these measures.

Pairwise p-values are Sidak-adjusted for the three group comparisons; no further adjustment across measures is applied.

Analyses are parametric throughout: one-way analysis of variance followed by Sidak post-hoc contrasts, which use the pooled within-group variance and adjust for the three pairwise comparisons; the pairwise p-values reported are adjusted. Variances were unequal for search time, number of fixations, P1, theta ITC and relative alpha power (Levene p=.002, .011, .010, .034 and .043). For P1 in SZ vs ASD, the unadjusted Welch t-test is reported alongside.

The dependence of the fixation-related response on saccade amplitude was tested with a linear mixed model of single-fixation P1 amplitude, with the amplitude of the preceding saccade as a covariate, group as a fixed factor, and participant as a random effect on both intercept and slope. Saccade amplitude was mean-centered, so that the group effect was estimated at the average amplitude. Fixed effects were tested with Wald chi-square tests. Separately, a linear regression of single-fixation P1 on the amplitude of the preceding saccade was fitted within each participant (86 participants); its value at a common amplitude of 6 degrees and its slope were compared between groups by one-way analysis of variance.

Subcortical-cortical and Cross-modal correlations were assessed using both linear regression (uncorrected for group) and ANCOVA, with group as a factor and fMRI measures used as covariates. The model included the group × covariate interaction. In ANCOVA, the proportion of residual error explained was calculated using the partial η² statistic. The cross-modal analyses were restricted to participants with usable search-task EEG (n=32; NC 5, ASD 16, SZ 11). Because of small sample sizes, cross-modal correlations were computed across the sample as a whole and were not estimated within group, except for the exploratory relationship between pulvinar activation and time to peak alpha suppression, which was tested as a group × covariate interaction.

Because the three modalities were acquired in overlapping but not identical samples, the sample size is stated with each result: 91 participants contributed to the primary behavioral analysis, 90 to the search-task EEG, 71 to the search-task fMRI and 32 to cross-modal (EEG/fMRI).

For all analyses, the pre-specified threshold for significance was p<.05. Data in the text are mean±sd unless otherwise specified.

## 3 Results

### 3.1 Experiment 1

A schematic of Experiment 1 is shown in Figure 1B-E

**Figure 1.**
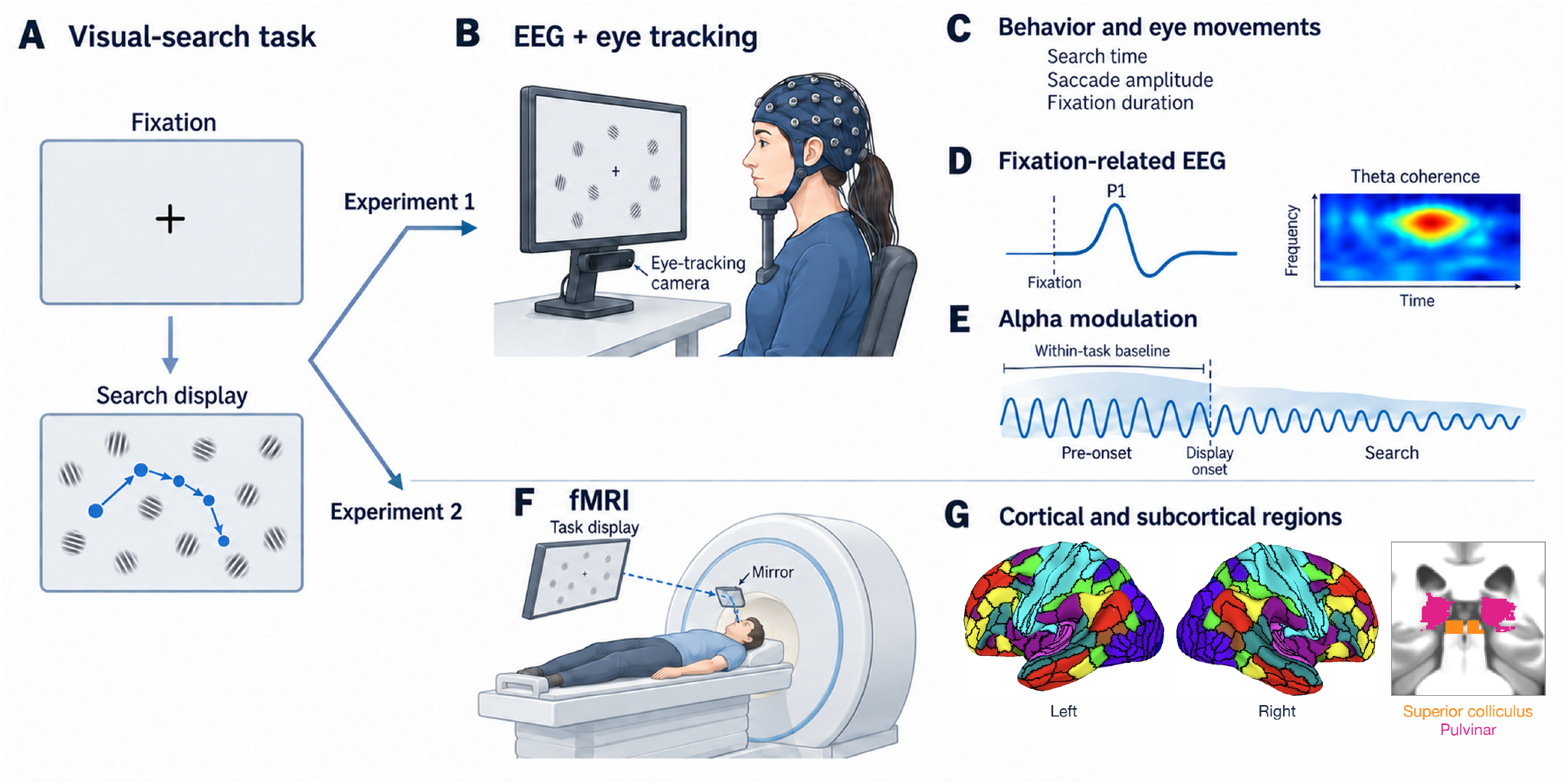
Schematic overview of the study. (A) Participants performed a guided visual-search task. (B) In experiment 1, continuous eye tracking and EEG were recorded during task performance. (C) Behavioral and eye-movement measures included search time, saccade amplitude, and fixation duration. (D) Fixation-related EEG measures included P1 amplitude, averaged from 80 to 140 ms at O1/Oz/O2, and theta intertrial coherence. (E) Posterior alpha power was measured relative to display onset, using the within-task pre-onset period as baseline. (F) In experiment 2, participants performed the task during fMRI, viewing the display through a mirror. (G) Cortical networks of Ji et al. (2019) and the superior colliculus and pulvinar masks of McFadyen et al. (2019) used in the fMRI analyses. Illustrations and waveforms in A–F are schematic; no participant data are plotted.

#### 3.1.1 Behavior

Mean search time differed significantly across the 3 groups (F(2,88)=9.06, p=.0003) (Figure 2A). In pairwise analyses, SZ showed significantly longer search times relative to NC (Sidak p=.003, d=0.74). Among autistic participants, search times were numerically shorter than in NC, a small-to-moderate difference (d=0.35) that is consistent with prior studies but not significant at this sample size. Nevertheless, the difference versus SZ was statistically reliable (Sidak p<.001, d=0.93).

**Figure 2.**
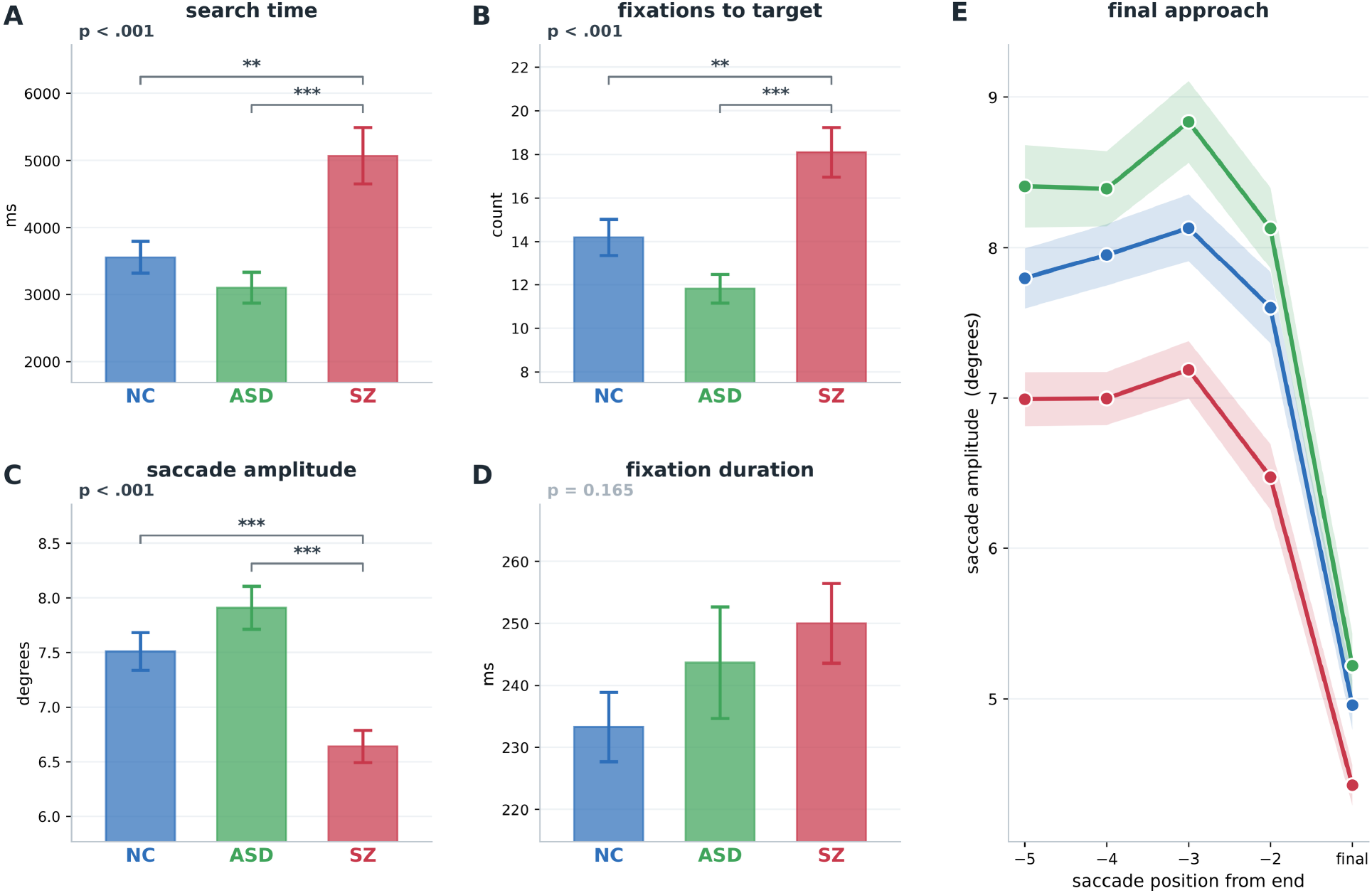
Mean ± SEM behavioral measures per search field. (A) Search time. (B) Number of fixations required to reach the target. (C) Saccade amplitude. (D) Fixation duration, excluding the final (target) fixation. (E) Saccade amplitude by ordinal position from the end of the trial. The value at the upper left of each bar panel is the omnibus one-way ANOVA; brackets mark Sidak post-hoc contrasts. *p<.05; **p<.01; *** p<.001

#### 3.1.2 Eye-tracking

In order to assess the basis for the difference in overall search time, we examined fixation number and fixation duration separately. Across groups, SZ participants needed significantly more fixations per search field to reach the target (Figure 2B), which in turn was related to a significant reduction in mean saccade amplitude averaged across fixations and search field (Figure 2C). These measures were strongly correlated across groups (r=-0.54, p<.0001). In contrast, fixation duration was not significantly different across groups (Figure 2D). Also, a control analysis was performed relative to the final approach (Figure 2E). Saccade amplitude was reduced in SZ at every ordinal position, including the saccade immediately preceding target fixation.

#### 3.1.3 Fixation-related potential (FRP)

Peak latency of the FRP-P1 (lambda) potential [Dimigen & Ehinger, 2021] was equivalent across groups (NC 108.4 ± 9.6, ASD 108.4 ± 10.0, SZ 108.4 ± 15.9 ms; F(2,83)<0.01, p>.99), leading to a common analysis window (Figure 3A). As expected, the P1 amplitude was significantly different across groups (F(2,83)=11.65, p<.0001) (Figure 3A,B). Pairwise testing showed a significant reduction in SZ relative to NC (Sidak p<.001, d=1.15); the reduction relative to autistic participants was large but not significant after correction (Sidak p=.067, d=0.88; unadjusted Welch p=.009, reported as a sensitivity test because variances were unequal); NC and autistic participants did not differ (Sidak p=.22, d=0.44). Topographies showed the same posterior positivity in all groups on a common scale (Figure 3C). Theta ITC at fixation onset showed a highly significant reduction in SZ relative to both NC and autistic participants, and no significant difference between autistic participants and NC (F(2,82)=12.85, p<.0001; SZ vs NC Sidak p<.001, d=1.21; SZ vs ASD Sidak p=.009, d=1.03; ASD vs NC Sidak p=.55, d=0.31) (Figure 3D, E). The number of accepted fixation epochs differed between groups (means NC 2285, ASD 1626, SZ 2945; F(2,83)=10.44, p=.0001), because SZ participants made more fixations, but it was unrelated to P1 amplitude overall (r=0.05, p=.68), within any group (all p≥.31) and after group-mean centering (r=0.11, p=.32).

**Figure 3.**
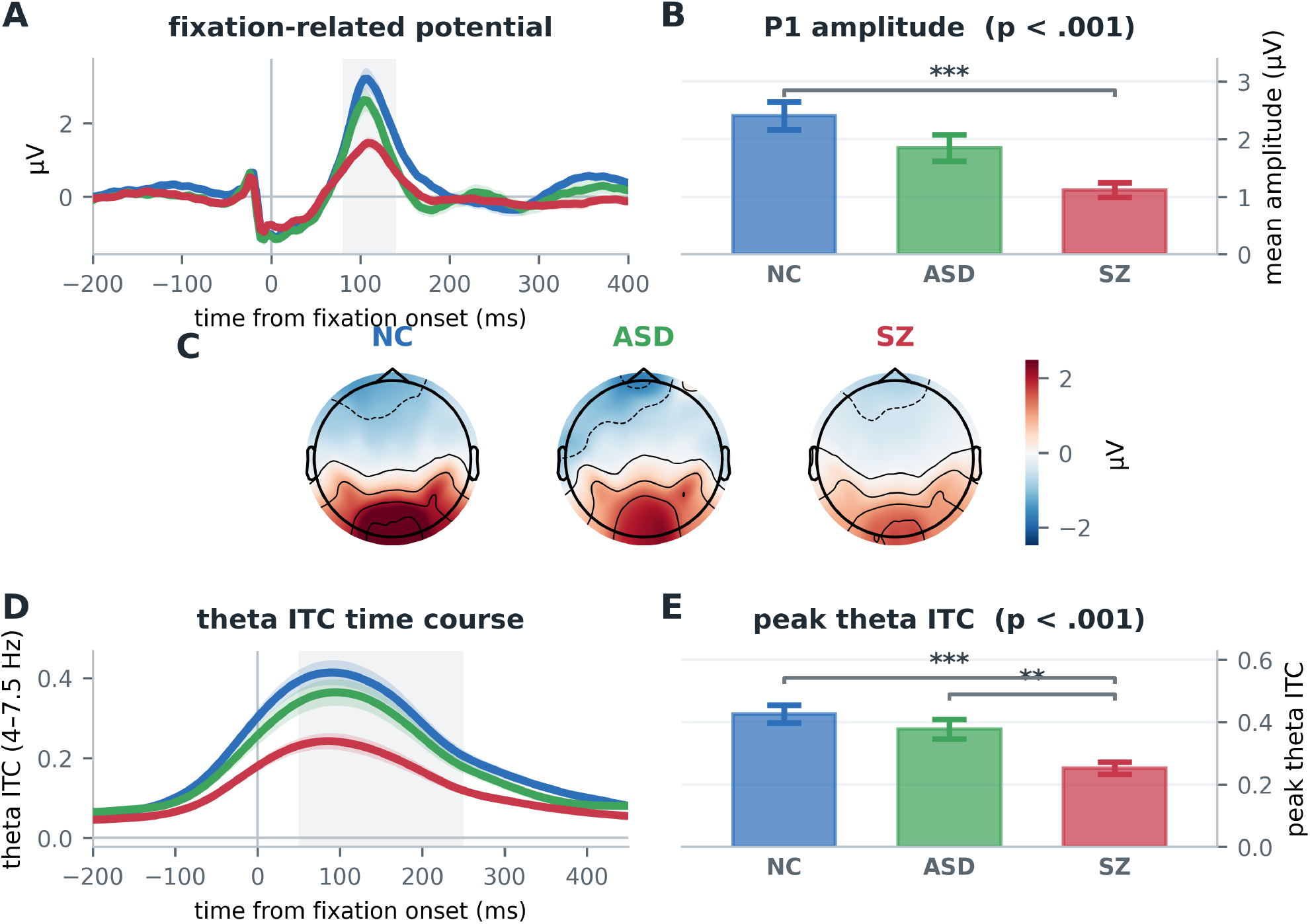
Fixation-related potential (FRP) measures (A) FRP over O1/Oz/O2 relative to fixation onset; shading is ± SEM, and the gray band marks the 80-140 ms analysis window. (B) Mean P1 amplitude by group (± SEM). (C) Scalp topographies over the 80-140 ms interval. (D) Fixation-locked theta ITC (4-7.5 Hz) over O1/Oz/O2; the gray band marks the 50-250 ms window. (E) Peak theta ITC by group (± SEM). In (B) and (E), the value at the upper left is the omnibus one-way ANOVA; brackets mark Sidak post-hoc contrasts *p<.05; **p<.01; ***p<.001.

Because mean saccade amplitude differed between groups (Figure 2C), we performed a control analysis to analyze FRP amplitude by saccade size within subjects for each group using a mixed-model regression (165,704 fixations from 85 participants; participant as a random effect on intercept and slope) (Figure 4).

**Figure 4.**
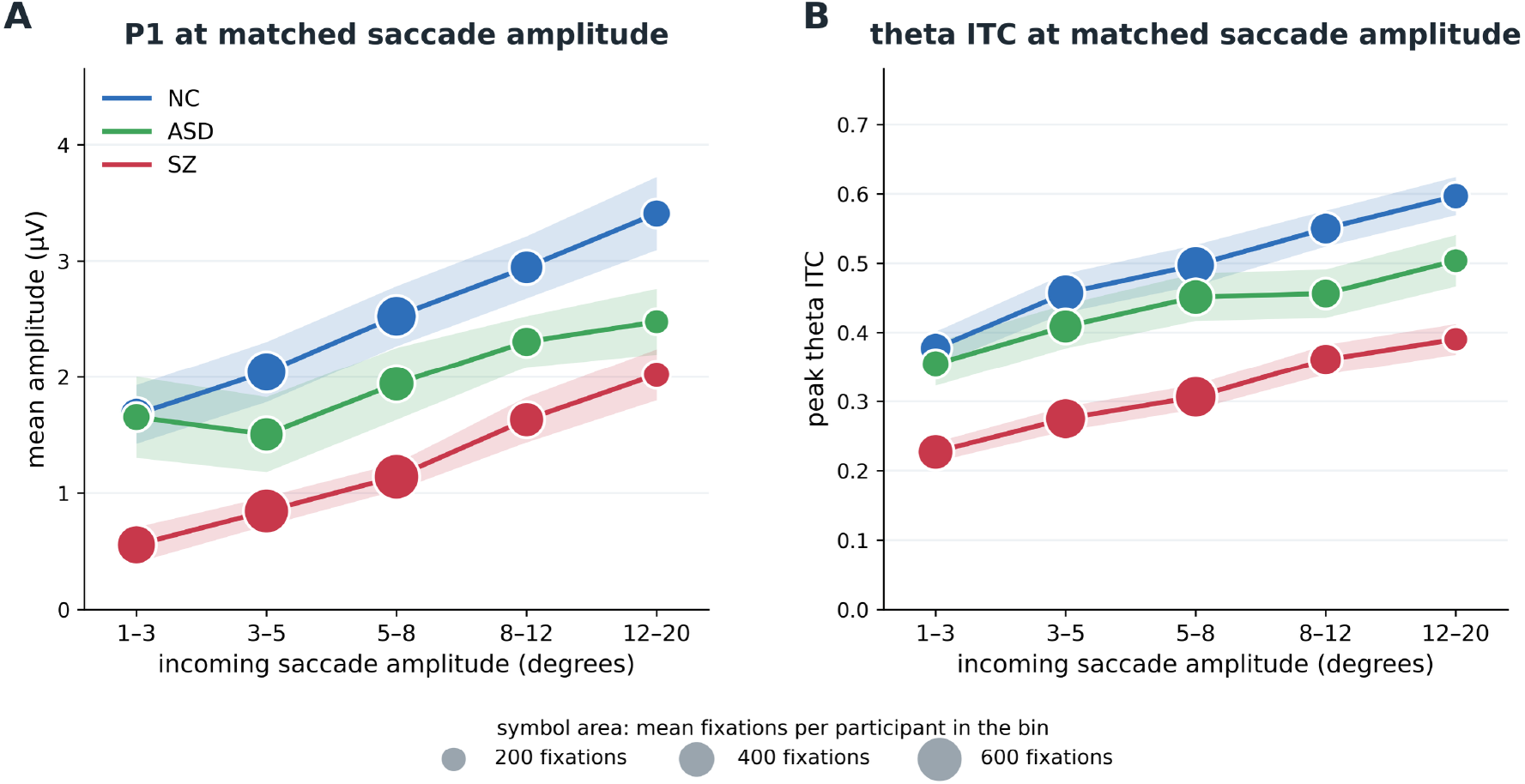
Analysis of P1 (A) and theta ITC (B) amplitude by saccade amplitude across groups. Lines are group means of participant bin means; shading is ± SEM. Symbol area is proportional to the mean number of fixations per participant in each bin (key); for theta ITC this is the number of fixations available, and ITC was computed on a fixed number of epochs per bin. P1: group F(2,81)=10.46, p<.001; group x bin F(8,324)=1.71, p=.094. Theta ITC: group F(2,70)=16.91, p<.001; group x bin F(8,280)=2.39, p=.017.

As predicted [Dimigen & Ehinger, 2021], FRP amplitude significantly increased with preceding saccade amplitude both across groups (χ²(1)=81.1, p<.0001), and within each group individually (NC χ²(1)=30.1, ASD χ²(1) = 7.8, SZ χ²(1)=7.0; all p≤.01). The slopes were similar across groups as reflected by a non-significant group × covariate interaction (χ²(2)=1.37, p=.50). The between-group difference remained significant following covariation for saccade amplitude (χ²(2)=20.7, p<.0001), as did the SZ vs. NC contrast (-1.27 µV, z=-4.54, p<.0001) (Figure 4A).

Theta ITC, evaluated at a common saccade amplitude of 6 degrees from a line fitted through each participant’s amplitude bins, including participants without all five bins, was likewise reduced in SZ (0.472, 0.419 and 0.293 in NC, ASD and SZ; NC 34, ASD 17, SZ 32; F(2,80)=16.54, p<.001), with SZ below both NC (Sidak p<.001, d=1.42) and ASD (Sidak p=.004, d=1.15). (Figure 4B)

##### Control analyses

A participant-level analysis of covariance on mean values gave the same P1 result as the mixed model (group F(2,81)=8.95, p=.0003; SZ vs NC p<.0001). The group × amplitude interaction was significant when participant was modeled with a random intercept only (χ²(2)=19.1, p<.0001) but not when the random slope was retained; the random-slope model is reported because slopes varied across participants. The group effect and the SZ vs NC contrast were unchanged under either specification.

Two further controls addressed residual confounding. First, the group difference did not vary across amplitude bins for P1 (group × bin F(8,324)=1.71, p=.094), so there was no evidence that the reduction was confined to particular saccade amplitudes; the group effect on the across-bin mean was F(2,81)=10.46, p<.001. For theta ITC the interaction was significant (F(8,280)=2.39, p=.017), with the group effect F(2,70)=16.91, p<.001. Residual differences in mean saccade amplitude within a bin were at most 0.17 degrees, which at the measured within-participant slope of 0.11 µV per degree could account for at most 1.3% of the group difference.

##### Relationship between FRP P1 and subsequent saccades

Although P1 amplitude was significantly predicted by the amplitude of the prior saccade, it did not significantly predict the size of the subsequent saccade (mean beta=-0.005; t=-1.47, df=84, p=.15). Theta ITC correlated weakly with search time (r=-0.23, p=.032) and P1 did not (r=-0.18, p=.097), and the theta ITC association did not survive group-mean centering (r=-0.05, p =.67).

Treating amplitude as continuous, in a regression fitted within each participant, gave the same result: at a common amplitude of 6 degrees, predicted P1 was reduced in SZ relative to NC (0.88 vs 2.12 µV, Sidak p<.001, d=1.08) and autistic participants (1.83 µV, Sidak p=.017, d=0.96), with no difference between NC and autistic participants (Sidak p=.76; F(2,83)=10.10, p=.0001). The d values of 1.08 and 0.96 use the unweighted average of the two group variances. The slope did not differ between groups (0.139, 0.081 and 0.101 µV per degree; F(2,83)=1.76, p=.18). Second, when the comparison was restricted to temporally isolated fixations, the SZ reduction remained significant at every threshold (NC vs SZ d=1.18, 1.11 and 1.07 at 200, 250 and 300 ms; all Sidak p<.001), and autistic participants did not differ from NC (d=0.43, 0.26 and 0.22; all Sidak p≥.22). Fixation durations were, if anything, longer in SZ (Figure 2D), so SZ participants had less overlap, which would inflate rather than reduce their P1.

Finally, P1 did not predict search time (r=-0.18, p=.097). The link between P1 and mean saccade amplitude (r=0.33, p=.002) is a difference between groups and did not survive group-mean centering (r=0.16, p=.13). P1 and theta ITC also correlated with the prefinal distance to target (P1 r=0.27, p=.014; ITC r=0.28, p=.008), replicating the previously reported relationship between fixation-related theta and parafoveal span.

#### 3.1.4 Alpha modulation

Ongoing alpha activity varied significantly across groups (F(2,76)=9.41, p=.0002) during the pre-search fixation intervals with significant elevation in the autistic group, relative to both NC and SZ (both Sidak p<.001) (Figure 5A,B). Locked to search display onset, all groups showed rapid alpha suppression sustained through the trial (Figure 5C). In order to compare suppression levels between groups, spectra were computed (Welch, a single 1 s Hann window per trial, zero-padded) over the 1-2 s interval following search onset, relative to the second before display onset. Suppression was maximal over the 8-12 Hz range (Figure 5D), with the most prominent changes over bilateral occipitoparietal regions (Figure 5E).

**Figure 5.**
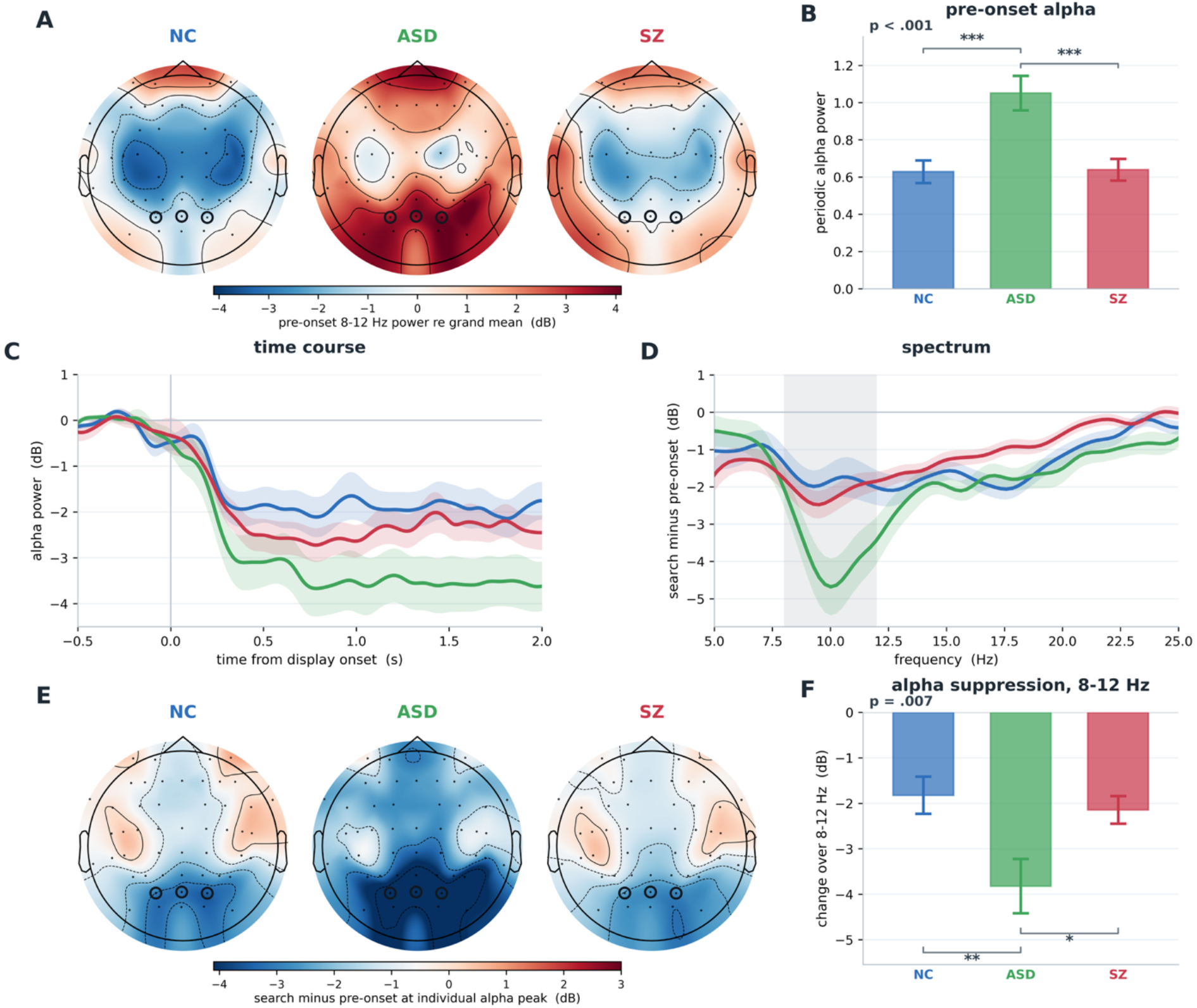
Alpha during visual search relative to pre-search baseline. (A) Pre-onset alpha power (8 to 12 Hz, -1000 to 0 ms relative to search display onset) for each group, in dB relative to the grand mean across channels and groups; rings mark PO3/POz/PO4. (B) Periodic pre-onset alpha power at PO3/POz/PO4, the height of the alpha peak above the aperiodic component (mean ± SEM; participants with a detected peak: NC 29, ASD 16, SZ 34). (C) Alpha power at PO3/POz/PO4 over time from display onset, relative to -0.6 to -0.2 s (mean ± SEM). (D) Change from the pre-onset window to the search window (1 to 2 s) across frequencies at PO3/POz/PO4. (E) Scalp distribution of the change from the pre-onset to the search window, read at each participant’s individual alpha peak frequency (participants with a detected peak); rings mark PO3/POz/PO4. (F) Participant-level mean change across 8 to 12 Hz, the band shaded in (D). Negative values indicate suppression. In panels B and F, the value at the upper left is the omnibus one-way ANOVA, and brackets mark Sidak post-hoc contrasts *** p<.001, ** p<.01, * p<.05.

Because the peak frequency was different across groups at baseline (F(2,76)=9.99, p=.0001), with SZ showing lower peak frequency relative to both NC (Sidak p<.001) and autistic participants (Sidak p=.006), the scalp distribution of suppression was measured at each individual’s peak alpha frequency (iAPF; Figure 5E). Suppression was summarized as each participant’s mean change across 8-12 Hz. That summary differed significantly across groups (F(2,83)=5.23, p=.007), with significantly greater suppression in ASD relative to both NC (Sidak p=.007) and SZ (Sidak p=.028) (Figure 5F).

Pre-onset alpha was higher in ASD than in NC and SZ, and the elevation was periodic, carried by the alpha peak above the aperiodic component (periodic alpha 1.051 vs 0.629 in NC, d=1.23; vs SZ d=1.17; both Sidak p<.001). The 8–12 Hz suppression was larger in ASD (d = .83, Sidak p = .007). It scaled with the pre-onset level (r = −.69); at matched pre-onset levels the group difference was d = .14, p = .66. Because that second is before the display, while the participant is already in the trial, these data do not separate a higher alpha level from greater modulation.

The same parameterization identified a second alpha abnormality, specific to SZ and involving frequency rather than power. Alpha peak frequency was slowed in SZ (10.51, 10.43 and 9.24 Hz in NC, ASD and SZ; F(2,76)=9.99, p=.0001), with SZ lower than both NC (d=-1.02, Sidak p<.001) and ASD (d=-0.98, Sidak p=.006), whereas autistic participants and NC did not differ (Sidak p=.99). Seven of 86 participants (NC 4, ASD 2, SZ 1) had no detectable alpha peak and did not contribute. The 8–12 Hz change is a fixed-band mean, so a shift in peak frequency could still contribute to it; the topography in Figure 5E is the change read at each participant’s own peak and is a separate measure. Slowed alpha peak frequency has previously been reported in SZ at rest [Ramsay et al., 2021; Murphy & Öngür, 2019; Yeum & Kang, 2018].

As opposed to amplitude of alpha suppression, the time to peak alpha suppression did not differ significantly between groups (F(2,83)=0.76, p=.47).

#### 3.1.5 Correlations with MCCB and ADOS-2

Clinical scores were correlated with the EEG measures within group. Correlations are Pearson and are not corrected for the number of tests.

In SZ, we observed a significant correlation between the FRP-P1 amplitude and the MCCB composite, such that lower P1 scores correlated with worse performance on the MCCB (r = .56, p = .001, n = 29) (Figure 6A). In exploratory analyses locked to fixation onset, the correlation with intertrial coherence was positive after onset in the theta and alpha range (Figure 6B), and the correlation with evoked power was positive in the alpha range over the same interval and negative at lower frequencies (Figure 6C).

**Figure 6.**
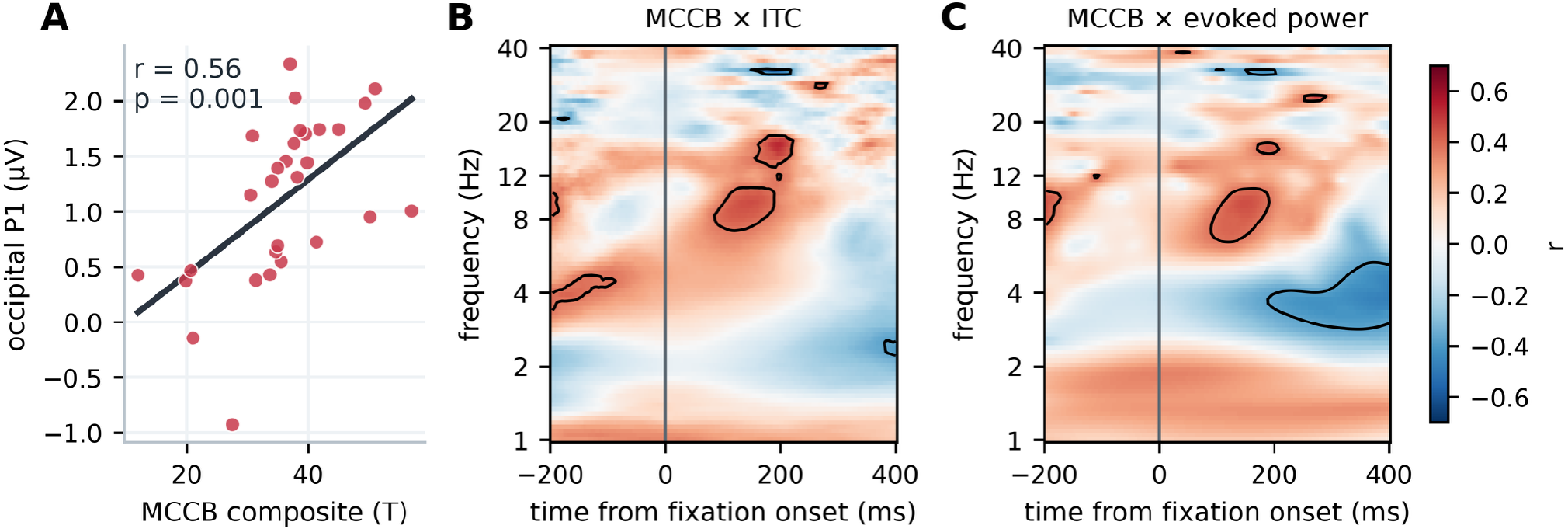
MCCB composite and the fixation-related response in SZ. (A) Occipital P1 against the MCCB composite (n = 29). (B) Correlation of the composite with intertrial coherence, locked to fixation onset (n = 28). (C) The same correlation for evoked power (n = 28). Contours mark uncorrected p < .05. Correlations are uncorrected. Two of the 31 participants with an MCCB composite (Table 1) had no P1; the maps use the 28 participants with a time–frequency measure.

Among the MCCB domains, P1 amplitude correlated with visual learning (r = .62, p < .001), working memory (r = .54, p = .002), attention/vigilance (r = .54, p = .002), verbal learning (r = .50, p = .005) and speed of processing (r = .47, p = .010), but not with reasoning/problem solving (r = .24, p = .21) (n = 30; 29 for speed of processing; Supplementary Figures S2–S6).

Across autistic individuals, we observed a significant correlation between FRP-P1 amplitude and the score on the ADOS-2 Communication scale. A higher score (greater symptoms) went with a larger P1 (r = .51, p = .024, n = 19) (Figure 7A). The ADOS-2 Communication score also correlated with the change in total 8–12 Hz power from the pre-onset second to the search window, such that a larger decrease in alpha correlated with more severe symptoms (r = −.51, p = .030, n = 18) (Figure 7B). The association with the power change was not confined to 8–12 Hz and was present throughout the β15–30 Hz range (Figure 7C). As opposed to the correlations with change score, absolute power during the search blocks did not correlate significantly with ADOS-2 Communication scores (r = .09, p = .74) (Figure 7D). ADOS-2 Social Interaction did not correlate with the 8–12 Hz change (r = −.29, p = .24, n = 18).

**Figure 7.**
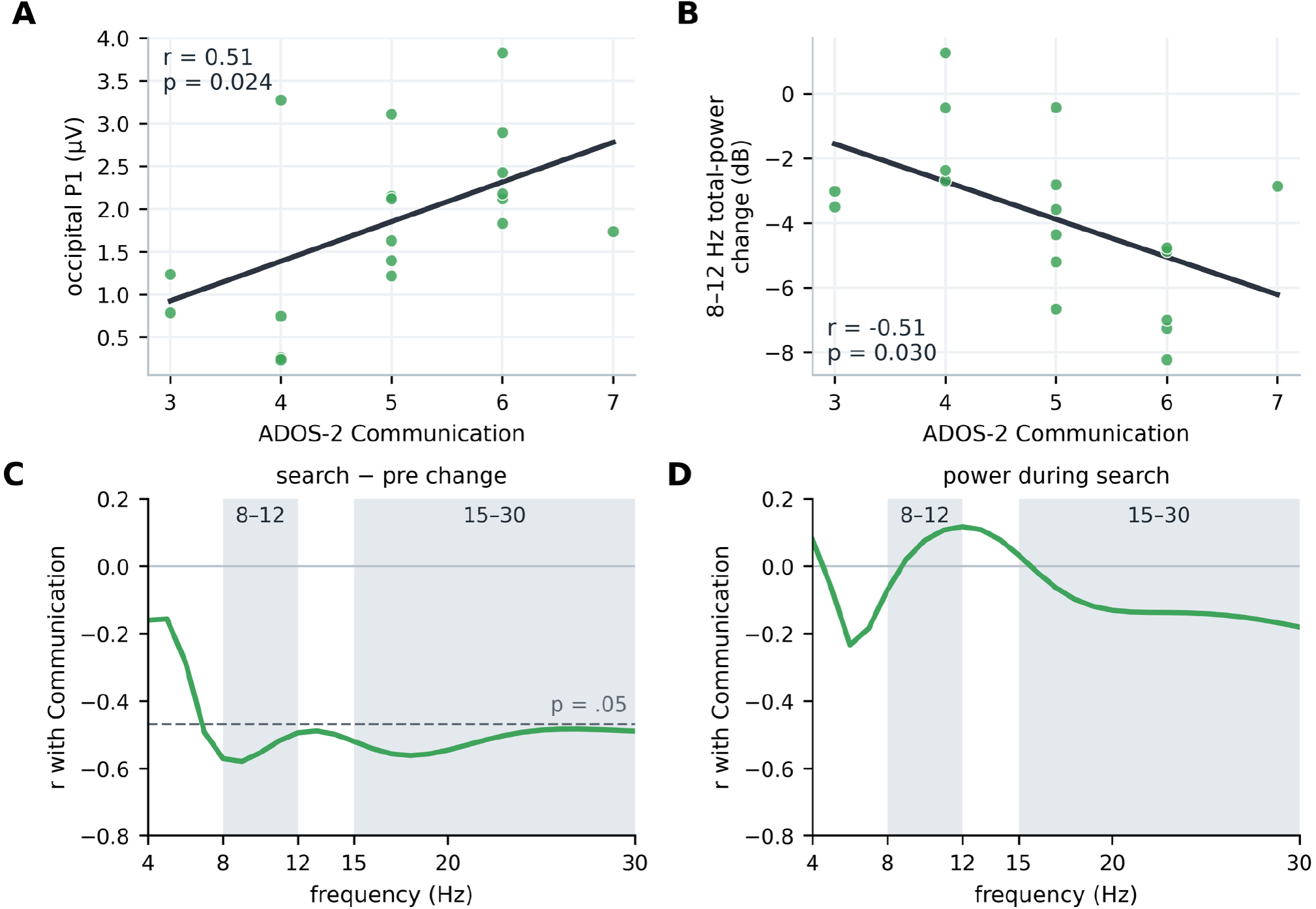
ADOS-2 Communication and the EEG measures in autistic participants. (A) Occipital P1 (n = 19). (B) Change in total 8–12 Hz power from the pre-onset second to the search window; more negative is a larger drop (n = 18). Panels A and B share 17 participants; A also includes two participants with no alpha spectrum, and B one participant with no P1. (C) Correlation of Communication with the search-minus-pre change at each frequency; the dashed line marks the r at which p = .05 (n = 18). (D) Correlation with power during search. The 8–12 Hz and 15–30 Hz bands are shaded. Correlations are uncorrected.

#### 3.1.6 Control analyses

With age or sex added as a covariate (one-way ANCOVA), every group effect reported above remained significant; sustained and peak suppression, which did not differ between groups, reached significance with age as a covariate (p = .020 and p = .040). Participants with SZ and NC were of similar age (35.5 and 35.3 years). Within SZ, antipsychotic dose (chlorpromazine [CPZ] equivalents, all 30 participants with a dose) did not correlate with P1 amplitude (r = −.16, p = .42) and correlated with theta ITC (r = −.39, p = .045); it did not correlate with search time, saccade amplitude or the alpha measures (|r| ≤ .22). Relative to NC, P1 and theta ITC were also reduced in participants at or below the median dose of 525 mg (P1 d = 0.82, p = .013; theta ITC d = 0.95, p = .004; n = 14; independent-samples t-tests). Education and estimated IQ, which were lower in SZ and are partly consequences of the illness, weakened the behavioral differences from NC (with education as a covariate, among participants with an education value, search time d = 0.71 to 0.36, p = .006 to .25; saccade amplitude p = .002 to .17) but not the EEG differences (P1, theta ITC and alpha peak frequency remained significant with either covariate, p ≤ .022).

### 3.2 fMRI

Primary fMRI analyses focused on the contrast between search and fixation blocks. Across groups, significant activations in search vs. fixation were observed in the Visual1 (paired t=8.5, df=70, p<.001), Visual2 (paired t=8.07, p<.001), Dorsal Attention (paired t=5.57, p<.001) and Frontoparietal (paired t=2.86, p=.006) networks (Figure 8A), whereas all other regions showed no significant difference between task conditions (all p>.3).

**Figure 8.**
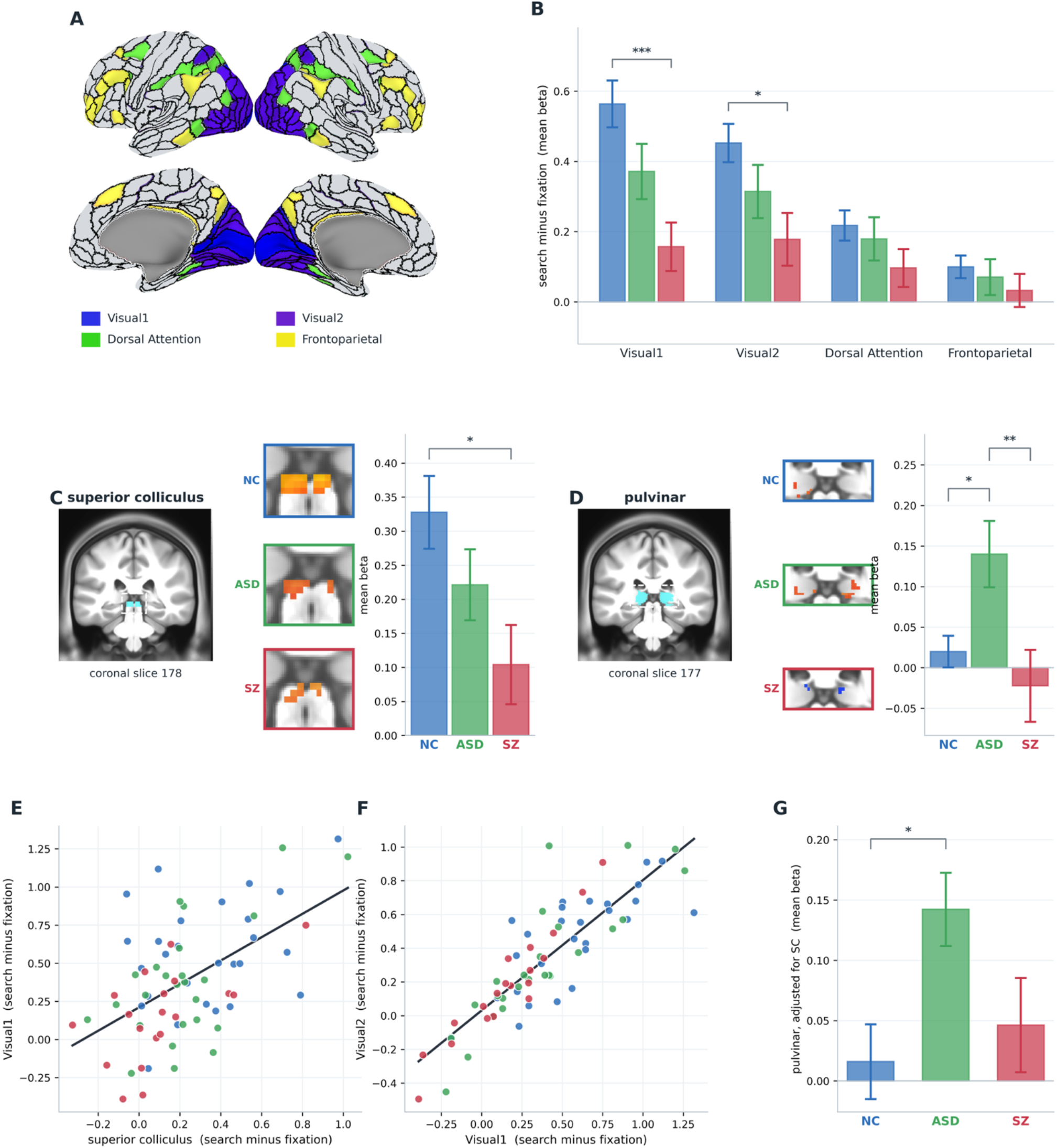
Cortical and subcortical modulation during search. (A) Mask showing the Visual1, Visual2, Dorsal Attention and Frontoparietal networks from [Ji et al., 2019]. (B) Mean search-minus-fixation beta for these networks by group. (C) Superior colliculus. *Left:* anatomical localization, showing location of the enlarged view from coronal slice 178 *Middle:* activation by group within the enlarged region. *Right:* mean ± SEM group activation values. (D) Pulvinar, as in (C), on coronal slice 177. (E) Collicular vs Visual1 modulation, one point per participant (r=.55, p<.001); the correlation was also significant within the ASD (r=.61, p=.001) and SZ (r=.59, p=.008) groups, as was the pulvinar-Visual1 correlation (r=.43, p=.033 and r=.53, p=.018). (F) Visual1 vs Visual2 modulation (r=.86, p<.001). (G) Pulvinar modulation by group adjusted for collicular modulation, from an analysis of covariance with a separate collicular slope for each group, because the group X collicular interaction was significant (F(2,65)=4.05, p=.022); estimated marginal means at the mean collicular value (± SE). Bars in (B)-(D) show means ± SEM. Brackets mark Sidak post-hoc contrasts. *p<.05; **p<.01; *** p<.001. Lines in (E) and (F) are least-squares fits across all participants.

Across the 4 regions, there was a highly significant group × task interaction (F(3,66)=26.2, p<.001), along with a significant main effect of group (F(2,68)=4.03, p=.022). Separate one-way ANOVAs were performed for each region. The main effect of group was significant for the Visual1 (F(2,68)=7.45, p=.001) and Visual2 (F(2,68)=3.86, p=.03), but not the Dorsal Attention (F(2,68)=1.24, p=.30) or Frontoparietal (F(2,68)=.56, p=.57) networks (**Table 2**). Both Visual1 (Sidak p<.001) and Visual2 (Sidak p=.022) differed significantly between the SZ and NC groups, whereas comparisons of autistic participants with NC and with SZ were all non-significant.

**Table 2.** Mean±std dev fMRI beta weights by network for task-related modulation (search minus fixation). F and p, one-way ANOVA across the three groups (p shown when below .05); post-hoc, Sidak contrasts with p below .05.

| **Network** | **NC (n=27)** |  | **ASD (n=25)** |  | **SZ (n=19)** |  | **ANOVA** |  | **Post-hoc** |
|---|---|---|---|---|---|---|---|---|---|
|  | **Mean** | **SD** | **Mean** | **SD** | **Mean** | **SD** | **F(2,68)** | **p** |  |
| **Visual1** | 0.564 | 0.347 | 0.372 | 0.393 | 0.157 | 0.301 | 7.45 | .001 | SZ<NC |
| **Visual2** | 0.453 | 0.284 | 0.314 | 0.379 | 0.178 | 0.328 | 3.86 | .026 | SZ<NC |
| **Somatomotor** | -0.050 | 0.151 | 0.016 | 0.217 | 0.023 | 0.135 | 1.30 |  |  |
| **Cingulo-Opercular** | 0.017 | 0.117 | 0.040 | 0.251 | -0.013 | 0.162 | 0.45 |  |  |
| **Dorsal Attention** | 0.218 | 0.224 | 0.179 | 0.307 | 0.096 | 0.235 | 1.24 |  |  |
| **Language** | -0.004 | 0.145 | 0.008 | 0.278 | -0.035 | 0.196 | 0.23 |  |  |
| **Frontoparietal** | 0.100 | 0.167 | 0.071 | 0.256 | 0.033 | 0.206 | 0.56 |  |  |
| **Auditory** | 0.018 | 0.199 | 0.020 | 0.300 | -0.017 | 0.190 | 0.16 |  |  |
| **Default** | -0.027 | 0.147 | -0.009 | 0.244 | -0.042 | 0.225 | 0.14 |  |  |
| **Posterior Multimodal** | 0.040 | 0.226 | -0.031 | 0.214 | -0.042 | 0.268 | 0.89 |  |  |
| **Ventral Multimodal** | 0.019 | 0.498 | -0.040 | 0.732 | 0.076 | 0.286 | 0.24 |  |  |
| **Orbito-Affective** | -0.043 | 0.251 | 0.142 | 0.404 | 0.011 | 0.209 | 2.48 |  |  |
| **Superior Colliculus** | 0.328 | 0.278 | 0.221 | 0.261 | 0.104 | 0.253 | 3.98 | .023 | SZ<NC |
| **Pulvinar** | 0.020 | 0.101 | 0.140 | 0.204 | -0.022 | 0.194 | 5.71 | .005 | ASD>NC, ASD>SZ |
| **Frontal Eye Fields** | 0.148 | 0.264 | 0.131 | 0.290 | 0.112 | 0.268 | 0.10 |  |  |

Finally, in order to relate the present values to recent post-mortem studies of Sz, values were extracted from cortical regions V1, V2, posterior parietal cortex and DLPFC as defined by Schoonover et al., 2024. Within each region, the MMP1.0 parcel showing the greatest cross-group activation was used for analysis. Significant activation deficits were observed in SZ for regions primary and secondary visual cortices and posterior parietal cortex relative to NC, but not in DLPFC. In contrast, no significant differences were observed for autistic participants relative to NC (Table 3).

**Table 3.** Mean±std dev fMRI beta weights by parcel for task-related modulation (search minus fixation).

| **Region (parcel)** | **NC (n=27)** |  | **ASD (n=25)** |  | **SZ (n=19)** |  | **ANOVA** |  | **Post-hoc** |
|---|---|---|---|---|---|---|---|---|---|
|  | **Mean** | **SD** | **Mean** | **SD** | **Mean** | **SD** | **F(2,68)** | **p** |  |
| **BA17 (V1)** | 0.771 | 0.436 | 0.524 | 0.476 | 0.306 | 0.500 | 5.61 | .006 | SZ<NC |
| **BA18 (V2)** | 0.559 | 0.373 | 0.416 | 0.385 | 0.242 | 0.404 | 3.76 | .028 | SZ<NC |
| **BA7 (7PL-R)** | 0.786 | 0.443 | 0.457 | 0.721 | 0.285 | 0.749 | 3.75 | .028 | SZ<NC |
| **BA46 (46)** | 0.069 | 0.224 | 0.082 | 0.279 | 0.032 | 0.213 | .24 | .80 |  |

#### Subcortical processing

In subcortical regions, significant task-related modulation was observed in SC (paired t=7.02, df=70, p<.001) (Figure 8C) and pulvinar (paired t=2.39, df=70, p=.02) (Figure 8D). In SC, there was a significant overall main effect of group (F(2,68)=3.98, p=.023) with a significant reduction in modulation in SZ relative to NC (Sidak p=.019), whereas autistic participants and NC were not significantly different. In correlational analysis, SC modulation correlated significantly with Visual1 network activation across groups (r=.55, n=71, p<.001) (Figure 8E). In order to parse the relative effects of SC activation vs. group, an ANCOVA was performed with group as a factor and SC activation as a covariate. In this analysis, the task-related modulation of SC activation significantly predicted modulation of Visual1 region activation following control for group (F(1,65)=21.3, p<.001), with no significant group × covariate interaction (F(2,65)=1.37, p=.26). The group difference in Visual1 modulation also remained significant (F(2,65)=3.93, p=.025) with a significant residual reduction in Visual1 modulation in SZ relative to NC (Sidak p=.028), suggesting additional local dysfunction at the level of the Visual1 network.

However, cortical activations were all highly correlated (r=.86, p<.001; Figure 8F) and none of the correlations remained significant following control for Visual1 activation, suggesting that interactions of SC were primarily with the Visual1 network.

In pulvinar, there was also a significant main effect of group (F(2,68)=5.71, p=.005), with significant enhancement of pulvinar modulation in autistic participants relative to NC (Sidak p=.038) and SZ (Sidak p=.007) (Figure 8G). Pulvinar modulation also correlated significantly with Visual1 modulation across groups (r=.33, p=.005). As with SC, the correlation remained significant in an ANCOVA analysis controlling for group (F(1,65)=5.15, p=.027) with no significant group × covariate interaction (F(2,65)=.35, p=.7). However, when the effects of SC and pulvinar were considered simultaneously relative to Visual1 modulation in an ANCOVA, the effects of SC remain strongly significant (F(1,66)=13.0, p<.001) whereas the effects of pulvinar were significant only at trend level (F(1,66)=3.38, p=.07).

#### Correlations between subcortical and cortical activation

SC activation correlated most strongly with activation in the Visual1 network (r=.55, p<.001). The correlation, moreover, remained strongly significant following covariation for group (F(1,65)=21.3, p<.001). The group difference in Visual1 activation remained significant, but at a much attenuated level (F(2,65)=3.93, p=.025). Overall, task-related SC activation accounted for 24.6% of the residual variance in Visual1 network activation, while group status accounted for an additional 10.8%. The group × covariate interaction was non-significant (F(2,65)=1.37, p=.26). Significant across-group correlations were also observed between SC and the Visual2 (r=.48, p<.001), Dorsal Attention (r=.48, p<.001) and Frontoparietal (r=.43, p<.001). However, given the covariation among networks, none remained significant following covariation for Visual1 network activation.

Pulvinar activation correlated most strongly with Frontoparietal network activation (r=.54, p<.001) (Table 4). Moreover, the correlation remained strongly significant (F(1,65)=20.0, p<.001) following covariation, whereas the residual group difference remained significant only at trend level (F(2,65)=3.03, p=.055). Overall, pulvinar activation accounted for 23.5% of the residual error in Frontoparietal network activation vs. 8.5% for group membership. The group × covariate interaction was non-significant (F(2,65)=.54, p=.58). Significant correlations were observed as well between Pulvinar and the DorsalAttention (r=.42, p<.001), Visual1 (r=.33, p=.005) and Visual2 (r=.27, p=.025) networks. However, given the covariation among networks, none of the correlations remained significant following covariation for Frontoparietal network activation.

**Table 4.** Correlations between Pulvinar Activation and indicated networks, controlled for group membership. Values are F(df1,df2). *Italic, p<.05; Bold Italic, p<.001*. Interaction effects were non-significant.

| **Factor** | **Pulvinar** |  | **Group** |  | **Group contrasts (Sidak)** |  |  |
|---|---|---|---|---|---|---|---|
| **Network** | **F(1,65)** | **p** | **F(2,65)** | **p** | **ASD-NC** | **SZ-NC** | **ASD-SZ** |
| **Visual1** | 5.15 | .027 | 8.22 | <.001 | .024 | .004 | .85 |
| **Visual2** | 2.93 | .092 | 0.27 | .76 |  |  |  |
| **Somatomotor** | 19.20 | <.001 | 3.58 | .033 |  |  |  |
| **Cingulo-Opercular** | 37.20 | <.001 | 3.95 | .024 |  |  |  |
| **Dorsal Attention** | 7.07 | .01 | 2.74 | .072 |  |  |  |
| **Language** | 31.70 | <.001 | 3.91 | .025 |  |  |  |
| **Frontoparietal** | 20.00 | <.001 | 3.04 | .055 |  |  |  |
| **Auditory** | 29.41 | <.001 | 4.38 | .016 |  |  |  |
| **Default** | 20.20 | <.001 | 1.44 | .25 |  |  |  |
| **Posterior Multimodal** | 19.20 | <.001 | 3.93 | .024 | .038 |  |  |
| **Ventral Multimodal** | 10.70 | .002 | 2.11 | .13 |  |  |  |
| **Orbito-Affective** | 11.30 | .001 | 0.48 | .62 |  |  |  |

In an exploratory analysis, we also regressed the SC activation patterns against cortex (Figure 9A). Consistent with the primary analysis, correlations fell primarily with vertices located within the Visual1, Visual2, DorsalAttention and Frontoparietal networks, with small, additional deactivations apparent in the Somatomotor network.

**Figure 9.**
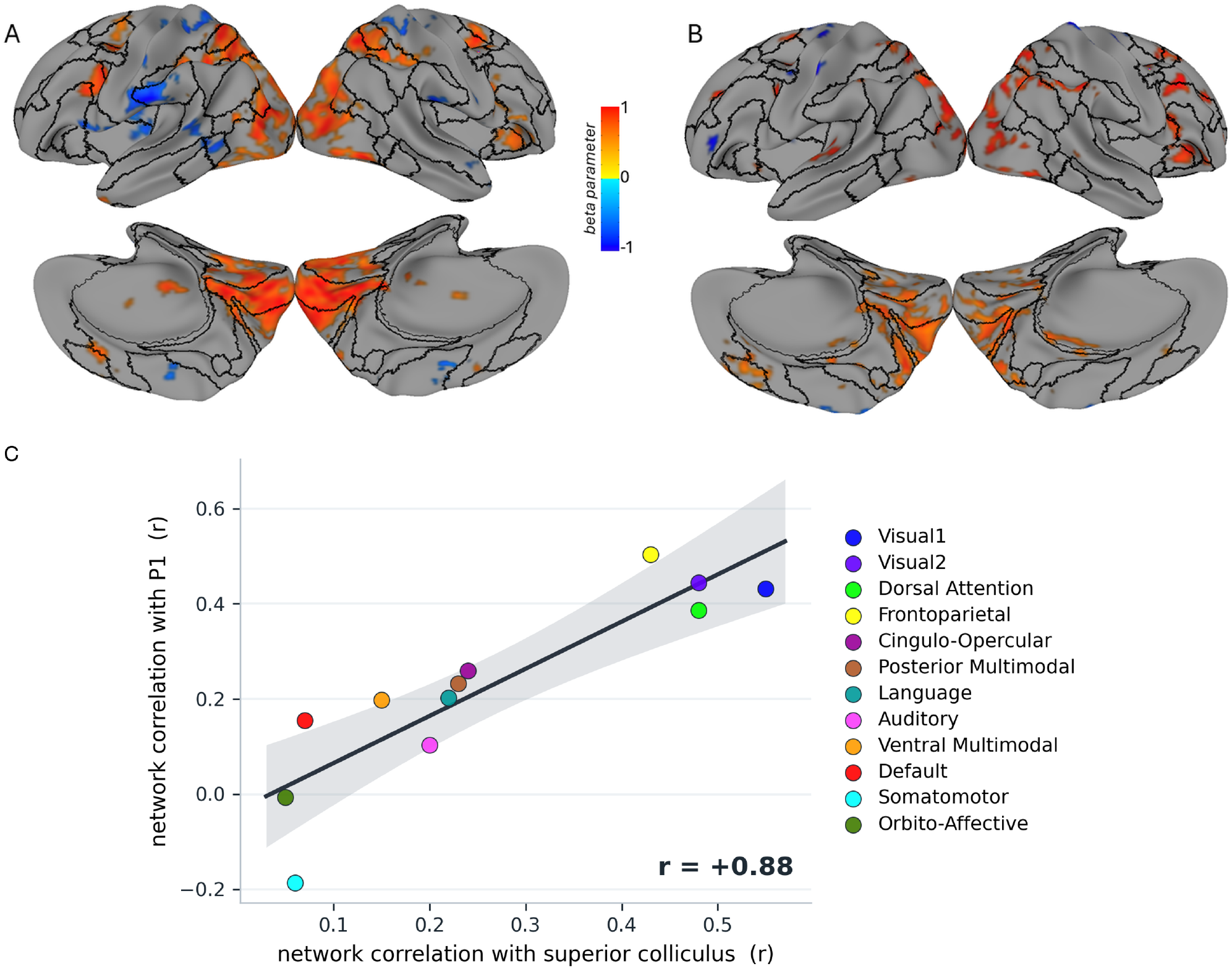
Cortex-wise correlation with (A) SC activation and (B) P1 amplitude. (C) Network profiles of the two correlations: for each of the 12 networks, the correlation of its search-minus-fixation activation with superior colliculus activation (n = 71) against its correlation with P1 amplitude (n = 32); r = 0.88. The same P1 profile against the pulvinar profile gives r = −0.40. The line is the least-squares fit and the band its 95% confidence interval.

#### FEF

In the network model used here, FEF is included within the Cingulo-Opercular network. Although the Cingulo-Opercular network as a whole did not show significant task activation, we conducted a separate analysis of the FEF given its potential importance to the search task. As predicted, significant bilateral FEF activation was observed across groups (t=4.11, df=70, p<.001). However, no significant between-group differences were observed (F(2,68)=.10, p=.9).

#### Analysis by condition

When fMRI data were analyzed separately for search and fixation blocks (Supplementary Table S1), significant between-group differences were observed for the Visual2 (F(2,68)=3.22, p=.046) and Dorsal Attention Network (F(2,68)=3.25, p=.045) in search blocks, and for the Visual1 (F(2,68)=5.99, p=.004) and Visual2 (F(2,68)=3.62, p=.032) in fixation blocks. In the search blocks, however, the pairwise differences between groups were non-significant (all Sidak p≥.056), whereas in the fixation blocks Visual1 activation was higher in SZ than in both NC (Sidak p=.005) and autistic participants (Sidak p=.023), and Visual2 modulation higher in SZ than in autistic participants (Sidak p=.028).

### 3.3 Cross-modal analyses

Cross-modal analyses were conducted to identify likely generators of the FRP P1. Across all participants with both modalities, P1 amplitude correlated most strongly with responses in the Frontoparietal (r=.50, n=32, p=.003) and Visual2 network (r=.44, p=.011) with somewhat weaker correlation to the Visual1 (r=.43, p=.014) and Dorsal Attention (r=.39, p=.029). However, in an ANCOVA, the correlations between P1 and Frontoparietal (F(1,26)=4.03, p=.055) and Visual2 (F(1,26)=3.22, p=.084) were significant at trend level only. In both cases, the group × covariate interaction was non-significant.

In an exploratory vertex-wide analysis, nominally significant correlations were observed primarily between P1 amplitude and vertices within the Visual2 network. (Figure 9B) Additional sites of correlation were present in the right Frontoparietal network. Overall, the pattern of activation between the two sets of correlation patterns were highly similar, with r = 0.88 across networks (Figure 9C). In contrast, the correlation of P1 vs. pulvinar activation (r = -0.40) was not significant, showing differential involvement across the two structures.

A significant cross-modal correlation was also observed between P1 amplitude in the EEG task and activation within both the Visual1 (p=.014) and Visual2 (p=.011) networks in the fMRI task (Figure 10). However, neither survives control for group membership. In contrast, correlations between P1 amplitude and FEF activation were non-significant both across and within-group.

**Figure 10.**
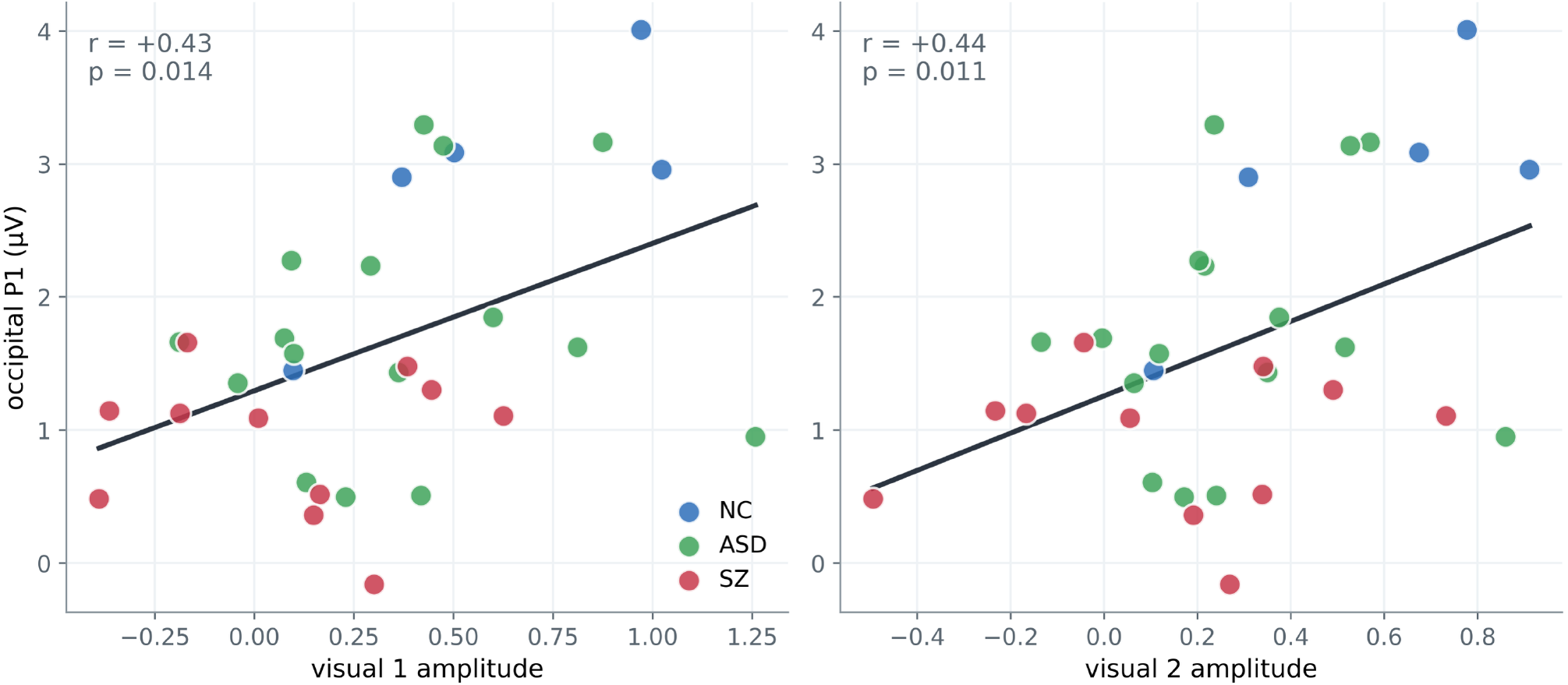
The occipital fixation-related P1 correlates with the visual-network BOLD response. Each point is one participant (blue, NC; green, ASD; red, SZ). (A) Occipital P1 amplitude vs the search-minus-fixation response of the Visual1 network (r=0.43, p=.014). (B) The same for the Visual2 network (r=0.44, p=.011). Lines are least-squares fits. Visual1 and Visual2 are adjacent, strongly correlated parcellations, so the two panels show one relationship rather than two independent results. The analysis uses the 32 participants with both usable search-task EEG and task fMRI (NC 5, ASD 16, SZ 11), a subset of the 71-participant imaging cohort. Neither correlation survives control for group membership (Results, cross-modal analyses).

Finally, analyses were conducted to evaluate cortical fMRI and EEG consequences of pulvinar hyperactivation in ASD. Increased pulvinar modulation was associated with increased activation most prominently with the Cingulo-opercular network (Figure 11A), but also across other networks related to cognitive and non-visual processing. The reduction in Visual1 activation in SZ remained strongly significant following covariation, with a somewhat weaker reduction in ASD. In addition, following covariation for the increase in pulvinar activity, ASD diagnosis was associated with a nominally significant decrease, rather than increase, in activity in the Posterior Multimodal network during search vs. fixation, although the analysis did not survive covariation for the number of networks tested.

**Figure 11.**
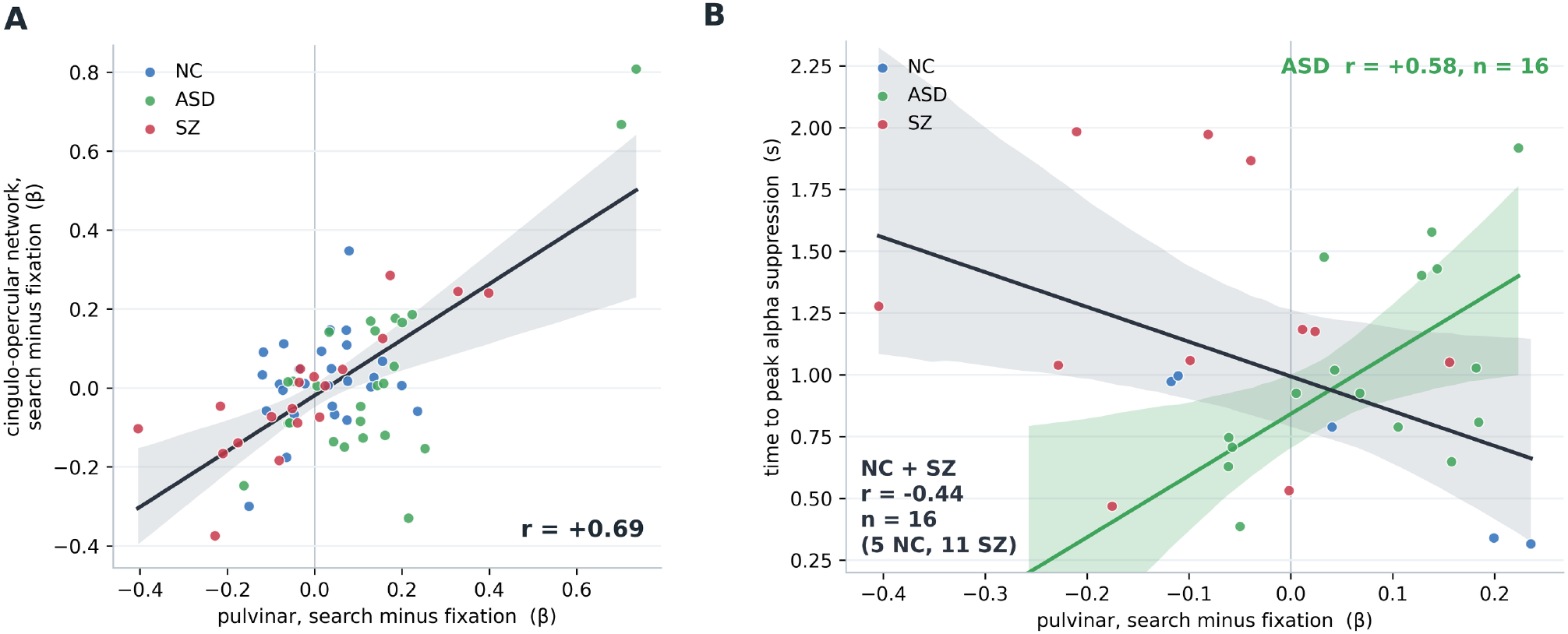
Pulvinar activation and its correlates. (A) Pulvinar vs cingulo-opercular network activation, search minus fixation, one point per participant (NC blue, ASD green, SZ red; n = 71: NC 27, ASD 25, SZ 19). The line is the fit across all groups with its bootstrap 95% band (r = 0.69). (B) Time to peak alpha suppression vs pulvinar activation in the participants with both EEG and fMRI (n=32: NC 5, ASD 16, SZ 11), fitted separately for ASD (r=0.58, p=.018) and for NC and SZ pooled (r=-0.44, p=.086), each with a bootstrap 95% band.

No significant correlations were observed between pulvinar activation and alpha-EEG measures across groups. Nevertheless, the correlation between pulvinar activation and time-to-peak alpha suppression correlated significantly within the autistic group (r=.58, p=.018), and in the opposite direction, but not significantly, in the NC and SZ groups pooled (r=-.44, p=.086). Moreover, the group × covariate interaction was significant when analyzed across all 3 groups (F(2,26)=3.99, p=.031) or between autistic participants and the combined NC and SZ group (F(1,28)=8.86, p=.006), indicating a differential slope across groups (Figure 11B).

## 4 Discussion

The guided search task has been extensively studied in neurocognitive research and is considered a paradigmatic test for evaluation of control of attention [Nuechterlein et al., 2009]. Deficits in search, as reflected in increased search times, have been extensively replicated in SZ [Dias et al., 2020; Nuechterlein et al., 2009; Sklar et al., 2020]. In contrast, ASD has been associated with normal or even superior search times [Kaldy et al., 2016; Constable et al., 2020]. In a previous study in SZ, we observed increased search times that were associated with reduced saccade size while performing the visual search, which, in turn, were associated with reduced fixation-P1 (lambda) responses. To our knowledge, this is the first study to evaluate FRP patterns in ASD in the guided search task, as well as the first study to evaluate fMRI activation patterns in either SZ or ASD.

The primary findings of the study are three-fold. First, SZ and ASD diagnoses were associated with the expected alterations in search times, with a significant, moderate-to-large increase in SZ (d=0.74) and a small decrease in search time among autistic participants (d=0.35) that did not reach statistical significance given the sample size. In SZ the change was accompanied by a reduction in mean saccade size, whereas saccade size in autistic participants did not differ from NC.

Second, SZ and autistic participants show differential patterns of alteration in EEG-based responses. Across all groups, P1 amplitude correlated strongly with the amplitude of the prior saccade (Figures 3, 4), supporting concepts that it is driven at least in part from corollary discharge from the oculomotor system [Rajkai et al., 2008]. Nevertheless, in SZ the P1 amplitude was reduced even following control for the amplitude of the incoming saccade, suggesting additional pathology at the level of visual cortex.

In the autistic group, P1 amplitude was not significantly different from NC; it was larger than in SZ, but not significantly after correction (Sidak p=.067, d=0.88; unadjusted Welch p=.009, a sensitivity test because variances were unequal). In contrast, among autistic participants, task-related modulation of the posterior alpha rhythm was significantly increased in magnitude, reflecting primarily higher alpha amplitude in the second before display onset (Figure 5), while in SZ the alpha modulation was similar to NC levels and significantly smaller than in ASD.

Third, in fMRI analyses, SZ and autistic participants showed differential alterations in activation patterns. SZ was associated with correlated reductions in SC, Visual1 and Visual2 network activation, but intact activation of the Dorsal Attention and Frontoparietal networks and of the frontal eye fields (F(2,68) = 0.10, p = .90) (Figure 8). Across groups, activation patterns related to SC activation were highly similar to those associated with P1 generation, supporting a potential role of SC in driving FRP P1 generation (Figure 9). In contrast, ASD diagnosis was associated with an isolated increase in pulvinar activation (Figure 8), that correlated strongly with the time to peak alpha suppression after search-display onset in the autistic group (Figure 11B).

Overall, these findings add to the growing literature demonstrating differential patterns of alteration in early visual alterations across SZ and ASD, such that SZ is characterized primarily by reduced activation within subcortical and early cortical visual regions, but relatively intact attentional engagement as reflected in either alpha modulation or activation of higher-tier brain networks including the DorsalAttention and FrontoParietal networks. Autism was associated with higher pulvinar activation and a higher alpha level before display onset. The 8–12 Hz suppression was larger among autistic participants (d = .83, Sidak p = .007). It scaled with the pre-onset level (r = −.69); at matched pre-onset levels the group difference was d = .14, p = .66. Because that second is before the display, while the participant is already in the trial, these data do not separate a higher alpha level from greater modulation. The study thus has specific neurobiological implications for both SZ and ASD, as well as mechanistic implications regarding FRP generation.

### 4.1 Schizophrenia (SZ)

Deficits in early visual processing in SZ were first reported in the early 1900’s by Busch and Kraepelin, who demonstrated that SZ was associated with an increased critical stimulus duration [Busch, 1904; Dondé et al., 2019]. However, sustained studies of visual processing dysfunction did not begin until the demonstration of visual backward masking deficits in the mid-1970’s [Saccuzzo & Braff, 1981]. More recent studies have demonstrated impaired contrast sensitivity and visual P1 generation that are indicative of reductions of non-linear gain preferentially within the subcortical magnocellular visual system [Martinez et al., 2008; Butler et al., 2005; Martinez et al., 2024]. Most recently, we have demonstrated impaired FRP generation [Dias et al., 2020; Dias et al., 2021], also indicative of dysfunction of early visual processing.

The present study further investigates the involvement of the early visual system using fMRI during the guided search task. As opposed to prior studies of visual processing in SZ that have primarily used stimuli presented at central fixation, the guided search paradigm involves repetitive saccadic eye movements, and thus additionally engages the Fronto Parietal (“task positive”) network, which is primarily involved in cognitive control; the Dorsal Attention system that is primarily involved in volitional control of eye movements; and the SC, which serves as a priority map and decision hub for implementing guided eye movements.

The primary fMRI finding is that activation of both the Fronto Parietal and Dorsal Attention networks were unaffected in SZ as was the FEF, whereas activation of the Visual1 and Visual2 networks and of the SC were all significantly reduced. Moreover, the region X group interaction was statistically reliable (p<.001), supporting the concept that impaired performance in this task was due primarily to dysfunction within low level sensory regions, rather than top-down control networks. The study thus supports similar findings within other cognitive control tasks, such as the AX-type visual continuous performance task [Dias et al., 2011]. In SZ, task-related modulation of the posterior alpha potential was also intact, further supporting preserved top-down attentional control mechanisms.

The study also sheds light on potential causes of FRP dysfunction in SZ. Based on scalp distribution, the FRP P1 generators localize primarily to visual sensory regions. The reduction in P1 amplitude is thus consistent with the observed reduction in Visual1 and Visual2 activation, which, in turn, may be driven at least in part by impaired SC input. The reduction in SC input into cortex may also contribute to the reduced saccade size observed in SZ, which in turn led to longer search times (Figure 2). Nevertheless, P1 amplitude was reduced even following control for saccade size (Figure 4), suggesting potential additional pathology at the level of visual sensory cortex.

The SZ correlation extends the findings from Dias et al. (2020). In that study, evoked theta power during guided search correlated, within SZ, with MCCB attention/vigilance (r = .56, p = .005) and processing speed (r = .50, p = .014). The association here is between the fixation-related P1 and the MCCB composite, not between evoked theta power and those two domain scores. The recordings also overlap with that study, so this is a further question asked of the same cohort, not an independent replication. What it adds is that cognitive performance tracks the early occipital response that is reduced in the group. The time-frequency maps show that the composite also tracks intertrial coherence and evoked power in the theta and alpha range after fixation onset. That is related to the earlier theta result. It is not the same measure.

### 4.2 Autism Spectrum Disorder (ASD)

Alterations in gaze patterns have been extensively documented in autistic individuals while processing social scenes. In contrast, performance on guided search with non-social stimuli is intact, or somewhat better than that of NC. In the present study, autistic participants were numerically faster than NC, but the difference was not significant, and their saccade amplitude did not differ. They showed elevated alpha power in the pre-search baseline. The larger task-related suppression followed that higher level, and pulvinar activation was increased. The FRP did not differ from that of NC. Schizophrenia shows the opposite pattern: a reduced FRP, with alpha modulation left intact.

The larger alpha suppression in the autistic group is consistent with the concept of hyper-engagement to non-social stimuli in ASD. However, it followed the higher pre-onset level (d = .14 at matched pre-onset levels), and within the group power during search did not correlate with ADOS-2 Communication (r = .09, p = .74), so these data do not separate greater engagement from a higher alpha level. The results of this study thus mirror findings from our recent studies of face processing in which increased stimulus-induced alpha suppression was also observed [Martinez et al., 2019; Martinez et al., 2021; Javitt et al., 2026].

The autism correlation is in the same direction as described by Martinez et al. (2019), who reported that a larger visual alpha suppression, measured at 7–14 Hz over 250–400 ms to simple visual stimuli, went with a higher ADOS Social Interaction score (r = −.62, p = .003). Similarly, Javitt et al. (2026) also reported a significant correlation between late alpha suppression during face processing and ADOS Social Interaction scores. In the present study, which measured responses to fixations rather than external stimuli, correlations are with the Communication score, whereas correlation with the Social Interaction score was not significant (r = −.29, p = .24). As opposed to the previous correlations, the correlation is also not confined to power in the alpha band. Thus, different aspects of alpha suppression abnormalities may contribute to differential symptom types in ASD.

The increase in pulvinar activation also mirrors the increases in pulvinar activation that we observed during motion [Martinez et al., 2019] or face [Martinez et al., 2021] processing in ASD. The pulvinar nucleus is known to play an important role in modulating cortical alpha activity [Saalmann et al., 2012]. In the present study, pulvinar activation did not correlate significantly with the amplitude of alpha modulation. Nevertheless, it correlated strongly with the time to peak alpha suppression, suggesting that it may play a critical role in the temporal evolution of alpha. Moreover, the direction of the correlation differed significantly between autistic participants and the other two groups, further supporting concepts of differential patterns of pulvinar function across SZ and ASD [Martinez et al., 2019, 2021]. Among autistic individuals, the differences were observed specifically in the periodic, rather than aperiodic component of alpha, further implicating thalamocortical feedback loops.

Consistent with theories of pulvinar function, pulvinar activation levels correlated strongly with activation levels across large areas of cortex. Once the increase in pulvinar activation was considered, ASD diagnosis was associated with nominally significant reductions in activation in both the Visual1 and Posterior Multimodal networks (**Table 4**). However, the results were exploratory and require independent confirmation.

### 4.3 FRP generation

In addition to providing insights into mechanisms by which subcortical and early cortical visual processing alterations contribute to alterations in cognitive performance across SZ and ASD, the present study adds information regarding neural mechanisms underlying FRP P1 generation. FRP P1 is widely used to evaluate visual processes in tasks associated with self-paced eye movements, including reading and naturalistic scene processing. At present, two main hypotheses have been proposed to account for the FRP P1 generation.

First, based upon the resemblance of FRP P1 (lambda potential) to the visual P1 elicited by external stimuli, it has been proposed that the FRP P1 represents input driven by the visual stimulus present at the point of fixation [Spiering & Dimigen, 2025]. Second, based on monkey intracranial recordings, it has been suggested that the FRP is driven by corollary discharge from the oculomotor system, leading to phase-reset of ongoing theta rhythms within cortex as a mechanism of increasing the efficiency with which the to-be fixated information may be processed [Rajkai et al., 2008]. In the monkey study, theta phase-reset was observed even in the dark. Moreover, the pattern of intracortical activation, characterized by activation of supra/infragranular layers of cortex rather than the thalamo-recipient layer, was consistent with the expected pattern of activation from corollary discharge inputs.

Here, because we used a uniform search field, the stimulation intensity at the point of fixation should be relatively invariant across saccade amplitudes, which should minimize afferent contributions to FRP scaling. We therefore evaluated two potential sources for the corollary discharge input into primary visual cortex: SC and FEF. In the case of SC, we observed a significant correlation between fMRI activation patterns related to SC activation and those related to P1 generation (Figure 9), supporting the concept that SC may serve as the source of corollary discharge input underlying FRP generation. P1 amplitude correlated with collicular search-minus-fixation activation (r = .40, n = 32, p = .023). The correlation did not survive correction for group membership (r = .09, p = .64).

In contrast, the correlation between P1 amplitude and FEF activation was not significant even in the across group analysis, arguing against FEF as a source of corollary discharge input into the SC, relative to P1 generation. Thus, the present finding support SC as a source of subcortical input that drives the phase reset of ongoing theta rhythms in visual cortex that, in turn, manifests as the FRP P1 response in time-domain analysis. Nevertheless, future simultaneous EEG/fMRI studies are needed to further analyze differential contributions of stimulus-driven vs. corollary discharge-driven activity in the initiation of the FRP P1 response.

### 4.4 Pathophysiological mechanisms

The present paper adds to the growing literature showing first that both SZ and ASD are associated with significant alterations in early visual processing and, second, that the patterns are differential across disorders. These findings also converge with recent post-mortem and gene expression studies.

Among autistic individuals, we observed a significant increase in pulvinar activation, consistent with our prior investigations, as well as the more general literature. Moreover, the relationship between pulvinar activation and alpha modulation was directionally opposite to that observed across the SZ and NC groups. These findings converge with post-mortem findings of increased pulvinar surface area in ASD and altered maturation both of pulvinar and peak alpha frequencies. In addition, the findings converge with recent findings that variations in gene expression patterns are most pronounced within the early visual system [Gandal et al., 2022].

In SZ, neurophysiological deficits in both auditory and visual function have been linked to disturbances in N-methyl-D-aspartate (NMDAR). These theories are supported by general findings of strong association between genetic alterations in the GRIN2A subunit of the NMDAR and risk for SZ, as well as by recent post-mortem studies showing reduced GRIN2A expression in visual and posterior parietal cortex in SZ, but not in DLPFC. Here, the pattern of fMRI activation deficit in SZ during the guided search mirrored the pattern of the GRIN2A reduction (**Table 3**). In SC, NMDAR play a critical role in multimodal integration and saccade initiation, raising the possibility that local NMDAR dysfunction within SC may contribute directly to the observed reductions in SC activation and FRP generation in SZ.

### 4.5 Limitations

Fixation-related responses were quantified by epoch averaging rather than by continuous-time regression, which deconvolves overlapping responses and models saccade amplitude and fixation duration within a single model of the unsegmented recording [Cornelissen et al., 2019]; overlap and saccade amplitude were addressed here by the isolated-fixation criterion, the amplitude-binned comparisons, and the trial-level mixed model instead.

No separate resting recording was obtained. The pre-onset measure is the second before the display appears, while the participant is already in the trial. The 8–12 Hz suppression was larger in autistic participants (d = .83, Sidak p = .007). At matched pre-onset levels the difference was d = .14, p = .66. These data do not separate a higher alpha level from greater modulation.

In fMRI, we used a group-average mask for whole pulvinar but did not collect tasks needed to further segregate into underlying nuclei. In a prior study, we observed that the increased pulvinar activity in autism is confined to inferior pulvinar, whereas reduced pulvinar activity in SZ is confined to lateral and medial divisions. Future studies need to further the role of specific pulvinar subnuclei in the different disorders.

Several findings were results of exploratory analyses, such as the differential correlation between pulvinar activation and time to peak alpha suppression between autistic participants and the other two groups and need to be confirmed in a priori investigations. Furthermore, the cross-modal correlations were limited both by the small number of individuals who participated in both experiments, as well as the fact that the EEG and fMRI were obtained in separate sessions.

## 5 Conclusions

In guided visual search, autism and SZ diverged from NC on different measures. In SZ, search was slower, saccades were shorter, and the fixation-related P1 and theta phase locking were reduced, while alpha modulation was preserved. In autism, search time and the fixation-related response did not differ from NC. Pre-onset posterior alpha power was elevated, and task-related suppression was also larger. Within each group, the fixation-related P1 correlated with a clinical score, the MCCB composite in SZ and ADOS-2 Communication in autism, and in autism a broadband drop in total power during search also correlated with Communication; these correlations were not corrected for multiple tests. These dissociable patterns support the use of fixation-related responses, together with posterior alpha, to separate early visual alterations across the two disorders.

## 7 Author Contributions

DCJ, EGM and AM conceived and designed the study and provided Project Administration and Supervision. EGM performed formal analysis and generated the original draft of the manuscript. AM, MAP, MG and AF contributed to data analysis. RT, ED, and OB contributed to patient recruitment and characterization. All authors contributed to review & editing of the final version.

## 9 Supplementary Material

Supplementary Results, Supplementary Methods, Supplementary Figures S1–S6 and Supplementary Table S1 are provided in the separate Supplementary Material file.

## 10 Data Availability Statement

The datasets analyzed for this study, together with the analysis code, are available from the corresponding author on reasonable request.

